# Global statistical constraint and local partner-sensitive inference support dynamic, open-ended verbal coordination

**DOI:** 10.64898/2026.10.03.756413

**Authors:** Tengfei Zhang, Daniel Joel Shaw, Siyuan Zhou, Lusha Zhu, Chunming Lu

**Affiliations:** State Key Laboratory of Cognitive Neuroscience and Learning & IDG/McGovern Institute for Brain Research, Beijing Normal University, Beijing, 100875, China; Behavioural and Social Neuroscience Research Group, Central European Institute of Technology (CEITEC), Masaryk University, Kamenice 5, Brno, 62500, Czech Republic; Department of Psychology, School of Life and Health Sciences, Aston University, Birmingham, B4 7ET, United Kingdom; Institute of Brain and Psychological Sciences, Sichuan Normal University, Chengdu 610066, China; School of Psychological and Cognitive Sciences and Beijing Key Laboratory of Behavior and Mental Health, Peking University, Beijing, China; IDG/McGovern Institute for Brain Research, Peking University, Beijing, China; Peking-Tsinghua Center for Life Sciences, Peking University, Beijing, China

**Keywords:** Open-ended communication, Verbal coordination, Deep language models, Error-driven updating, Pragmatic inference, fNIRS

## Abstract

Effective verbal communication requires individuals to align their communicative intentions through the precise use of utterances, yet how this is achieved in dynamic, open-ended communication remains unclear. Combining predictive deep language models (DLMs) with cognitive computational modeling, we tested a computational division of labor between context-sensitive global constraint on the utterance space and partner-sensitive local adjustment of selection priorities. We formalized local adjustment using an active-inference-inspired updating rule within a statistical– pragmatic hybrid model. Thirty dyads performed a dynamic, open-ended collaborative word-generation task during functional near-infrared spectroscopy (fNIRS) hyperscanning. Interlocutors converged in their lexical choices, and lexical alignment was best explained by a hybrid model integrating contextual selection priors with interaction-specific local adjustment among the models tested. This hybrid process formed a dynamic hypothesis landscape comprising a global semantic search region and local selection priorities. Critically, interaction did not simply reduce uncertainty: within individuals, the hypothesis space of global semantic search progressively narrowed while local uncertainty among plausible alternatives increased; between partners, both global hypothesis space and local selection became increasingly aligned, with alignment of local selection priorities most strongly predicting subsequent lexical alignment. The contribution of local adjustment was stronger during precise verbal communication than in the control tasks. Neural analyses further identified correlates of the local adjustment process: temporoparietal and anterior temporal regions supported partner-related inference and semantic prediction error, respectively, and their activity during thinking predicted subsequent inferior frontal activity associated with self-updating during speaking. The participant’s own hypothesis landscape was also represented in the left inferior parietal–supramarginal cortex. The main findings generalized across language models and an independent Czech dataset. Together, these findings reveal that alignment in open-ended communication emerges from complementary statistical and pragmatic computations, providing a framework for understanding human and human–AI communication.

## Introduction

Effective verbal communication depends on the ability of interlocutors to align their communicative intentions through precisely chosen utterances^1–3^. In this process, mutual understanding is gradually built and updated across turns^1,4–6^. Achieving such alignment remains a long-standing computational challenge during open-ended two-way communication, where speakers must select what to say from a vast and continually changing space of possible utterances. Unlike more constrained forms of language use, open-ended conversation does not provide a fixed communicative intention or a predefined set of candidate utterances in advance. Instead, speakers must generate and assess candidate utterances online while simultaneously tracking a partner whose evolving communicative intentions inform what the speaker intends to convey and how the speaker’s utterances are likely to be understood^7–10^. Although humans do this routinely in everyday conversation, it remains unclear how the human brain dynamically selects an appropriate utterance from an open-ended candidate space to achieve precise communicative alignment.

Two computational traditions are relevant to this problem. Deep language models (DLMs) formalize how preceding context narrows the space of likely upcoming linguistic units by generating probability-weighted distributions over candidate continuations, based on statistical regularities learned from naturalistic language^11,12^. This predictive generation capacity of DLMs parallels the context-sensitive predictive processing observed in the human brain during language comprehension and production^13–16^. Yet most DLMs acquire such abilities through large-scale text training rather than adaptive engagement in real-time, two-way communication like the human brain^12,16,17^. Although alignment training improves interactional performance^18^, DLMs still make systematic errors on tasks that require pragmatic prediction grounded in mental-state understanding^19,20^. Thus, DLMs might offer a useful approach to context-dependent utterance space compression, but still have limitations in determining how this space should be adjusted online according to an interacting partner’s evolving behaviour.

A second tradition comes from cognitive-computational models of pragmatic inference, which model communication as reasoning over why a speaker chose a particular utterance from a set of available alternatives. Related work has shown how speakers anticipate listeners’ interpretations and how listeners reason back from those utterance choices^8,10,21,22^. The active inference (AIF) framework further extends this perspective to ongoing interaction by treating communication as the continuous updating of partner-directed predictions within a hierarchical predictive architecture that supports uncertainty reduction and intentional alignment over time^23–26^. However, most existing models examined inference over predefined finite choice sets, leaving an open question of how to compress the utterance space over an open-ended communication context.

Taking both computational traditions together, we propose that achieving communicative alignment during an open-ended two-way communication depends on a hybrid computational mechanism that supports global-to-local utterance selection. Specifically, based on the evolving conversational context, context-dependent statistical prediction addresses the global search problem by compressing an initially open-ended utterance space into a contextually relevant semantic region. Based on prediction error, partner-sensitive inference then addresses the local problem of precise utterance selection by dynamically adjusting the priorities of candidates within the above region. The candidate utterances within this region, along with their partner-adjusted selection priorities, form a hypothesis landscape that may guide the open-ended communication process.

Similar hybrid computational approaches have appeared before. Some studies have incorporated DLMs into recursive pragmatic inference architectures to improve the communicative performance of artificial language systems^27–29^. In these models, DLM-derived probabilities of utterances or referents provide the basis for reasoning about speakers’ choices and listeners’ interpretations. In parallel, recent studies have utilized DLMs to supply components that are traditionally manually specified in cognitive pragmatic models (e.g., candidate utterances), thereby broadening their applicability to more open-ended communicative contexts^30–32^. In these approaches, DLMs generate and evaluate alternatives, while cognitive models specify the pragmatic computations over these alternatives. However, although these DLM–pragmatic hybrids offer valuable insights into integrating DLM-based statistical prediction with partner-sensitive pragmatic inference, how this computational division of labor unfolds across global and local scales during online, open-ended, two-way communication remains unclear.

In our hypothesized hybrid computational process, a central question, therefore, is how the global and local components of each interlocutor’s hypothesis landscape are progressively reshaped over interaction and, in turn, guide subsequent lexical selection. Here, the global component is operationally defined as the contextually compressed semantic region spanned by candidate utterances, and the local component as their partner-sensitive selection priorities. Previous work has shown that stronger or accumulating contextual information is associated with lower semantic dispersion among predicted lexical candidates and reduced uncertainty in the predicted semantic distribution of upcoming words^33,34^. Thus, we predict that, during the evolution of two-way communication, the accumulation of conversational context allows context-dependent statistical prediction to progressively compress each interlocutor’s candidate utterances within an increasingly specific semantic region, thereby reducing the global search stress. However, such global semantic compression may not resolve the local selection problem within the semantic region. Linguistic theories propose that semantically related lexical items can be co-activated and compete with one another during word selection, with greater lexical similarity giving rise to stronger competition among candidate words^35–38^. We therefore predict that, as the global semantic region becomes increasingly compressed, local selection uncertainty among the remaining plausible alternatives increases.

Another central question is whether and how the hypothesis landscape is neurally represented in the interlocutors’ brain. Evidence indicates that a latent high-dimensional semantic space derived from DLMs maps onto neural responses, and there is shared geometric structure between model-derived contextual embeddings and brain embeddings in classic language regions such as the inferior frontal gyrus (IFG)^13,16,39^. Moreover, given the reciprocal nature of two-way communication, such representations may become coordinated across interlocutors^40–42^, allowing the speaker’s hypothesis landscape to be reflected both in her/himself and the partner’s brain. Accordingly, we predict that the hypothesis landscape is represented in the brains of the speaker, the partner, or both.

To test the hybrid hypothesis and these questions, we recruited 60 participants (30 same-sex dyads) to perform a collaborative word-generation task adapted from prior work on dynamic verbal coordination^43^ (Fig.1). The task provided a tractable test of open-ended utterance selection because participants had to converge on the same word across repeated trials based only on each other’s prior responses. Here,”open-ended” refers to the absence of a predefined candidate set, rather than unconstrained natural conversation. Two control tasks dissociated precise lexical alignment from broader semantic coordination and from language production without interpersonal exchange (see Methods). Across tasks, we combined behavioral analysis, computational modeling, and functional near-infrared spectroscopy (fNIRS) hyperscanning, a method well suited to spoken interaction because it tolerates movement and operates silently ^44^. Behaviorally, we found that progressive semantic convergence emerged selectively during precise verbal collaboration. Computationally, the hybrid model combining context-dependent statistical prediction with error-driven, partner-directed inference best accounted for the emergence of semantic alignment. This model characterized a dynamic hypothesis landscape comprising a global semantic search region and local selection probability distribution over candidate words. Within individuals, the semantic search region progressively narrowed, while local selection uncertainty increased; between partners, both semantic search regions and local selection probability distributions became increasingly aligned. Neural analyses provided evidence consistent with a hierarchical organization of partner-directed updating and identified a neural representation of the participant’s own hypothesis landscape.

**Fig. 1.**
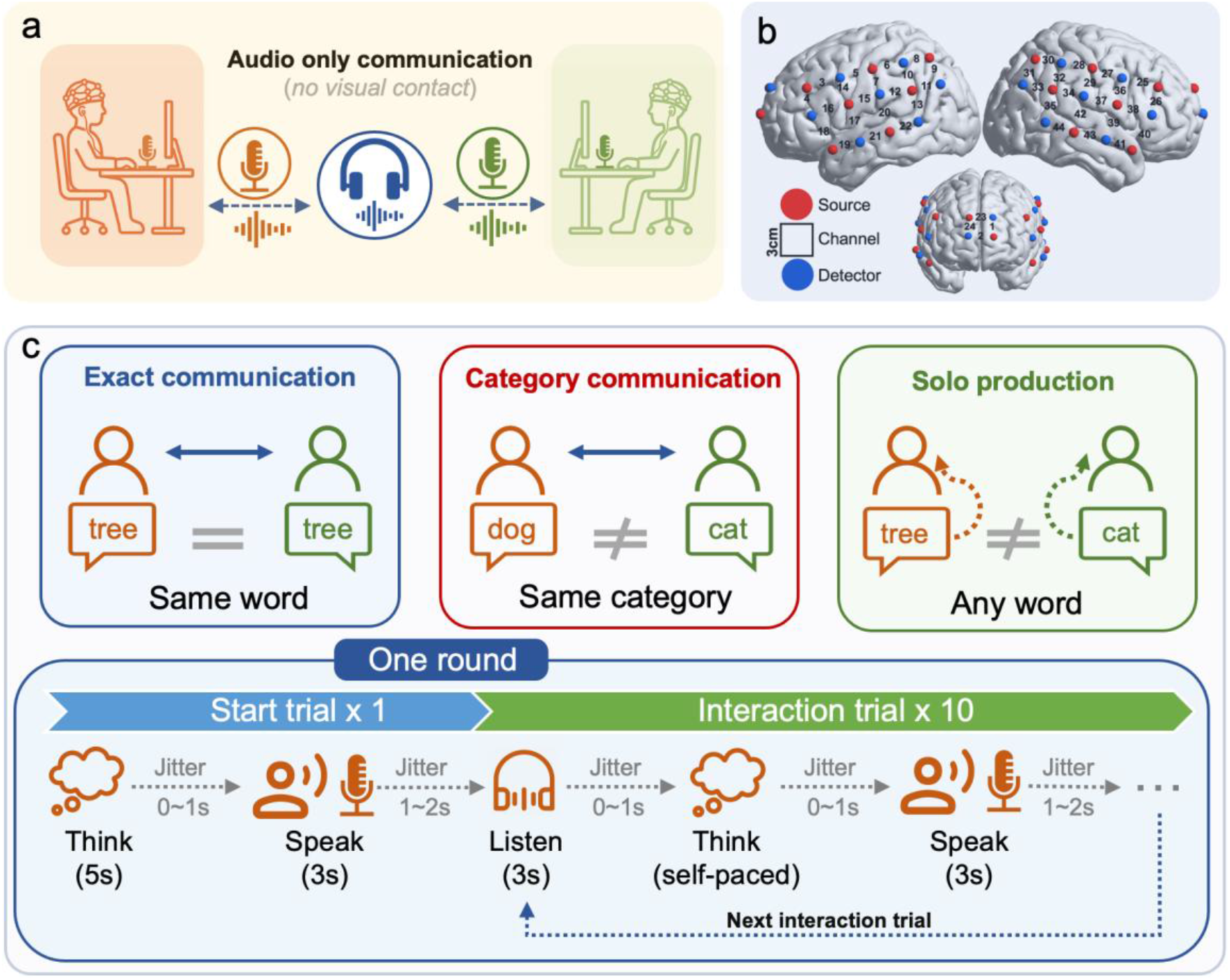
(a) Experimental setup. (b) Layout of fNIRS channels. (c) Overall design of the experiment: Each round consisted of one start trial without audio playback, followed by 10 interaction trials that involved listening, self-paced thinking, and speaking. In the *exact* or *category communication* tasks, participants heard their partner’s preceding word, whereas in *solo production* task they heard their own. Jitter intervals within each trial—between listening and thinking and between thinking and speaking—were independently sampled from a uniform distribution of 0–1 s. The between-trial interval, from speaking to the listening phase of the subsequent trial, was sampled from a uniform distribution of 1–2 s. Consecutive rounds were separated by 15 s.

Together, the results support the proposed statistical–pragmatic hybrid hypothesis.

## Results

### The exact communication elicited progressive semantic convergence

We first asked whether lexical alignment emerged progressively over the course of interaction (Fig. 2a and b). Here the alignment was quantified as the trial-wise semantic similarity between the two words produced by each dyad (see Methods). The Linear Mixed-Effects (LME) analysis, with the speaking and thinking durations as covariates, demonstrated a progressive pattern of the alignment over trials in the *Exact communication* task (*β* = 0.216, *t*(818) = 13.034, *p* < 0.001, *d* = 2.42; Fig. 2b), but no such pattern was observed in the *Category communication* and *Solo production* tasks (Details in Supplementary Text 2; the reaction-time analyses are reported in Supplementary Text 1).

**Fig. 2.**
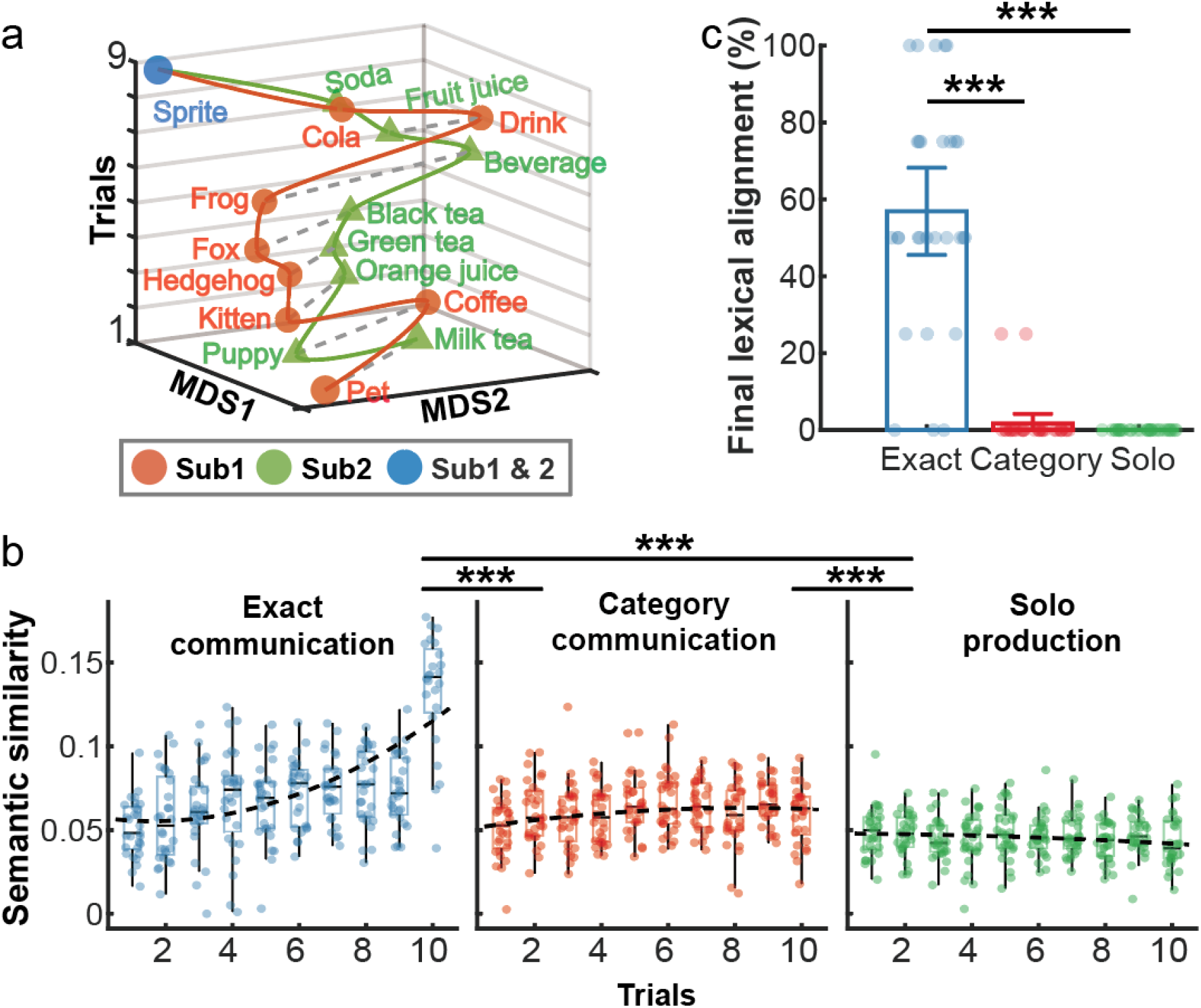
(a) An example word chain from a dyad during an *Exact communication* round. (b) The dynamic change of semantic similarity across trials in all three tasks. The first trial and repeated-word trials were excluded for each round in the *Exact communication* task. The remaining trials were aligned by their relative position, ensuring that each round contained the same number of consecutive interaction trials (10 trials) (see Methods). In box plots, the center line indicates the median, the top and bottom boxes correspond to the third and first quartiles, respectively, and the whiskers extend to the most extreme values within 1.5 times the interquartile range. (c) Final lexical alignment across tasks. A round was considered to reach final lexical alignment when the dyad produced the same word and continued to repeat it for the remainder of the round. Shown is the percentage of such rounds in each task.

We next examined whether this progressive pattern resulted in final lexical alignment. A round was considered to reach final lexical alignment when the dyad produced the same word and continued to repeat that word for all remaining trials of the round. In the *Exact communication* task, more than half of the rounds reached exact lexical alignment (56.9% of the 116 rounds, 29 dyads × 4 rounds; Fig. 2c), but such alignment was rare in the two control tasks (Category: 1.72%; Solo: 0%). An LME analysis further showed that the proportion of rounds showing exact lexical alignment was significantly higher in *Exact communication* than in either control task, while the two control tasks did not differ significantly from each other (See Table S4 for statistical details).

Finally, we tested whether the progressive pattern of the semantic alignment depended on whether final lexical alignment was achieved, and whether there was a practice effect across rounds. The results showed that the alignment increased significantly across trials in both successful (i.e. rounds showing final lexical alignment) and failed rounds (i.e. rounds not showing final lexical alignment, Fig. S3a, see Supplementary Text 2 for details). Moreover, semantic alignment remained stable across rounds, with no evident practice effect (Fig. S4a and Table S4, see Supplementary Text 2 for details). Together, these findings show that progressive lexical alignment is a robust and task-specific feature of *Exact communication*. We next asked what computations could account for how such convergence emerges from an initially open response space.

### Reconstructing the evolving lexical hypothesis landscape using a statistical– pragmatic model

#### Model overview

To reconstruct the evolving lexical hypothesis landscape underlying interaction, we developed a Statistical–Pragmatic Hybrid (SPH) model (Fig. 3a, see Methods SPH model). The model assumes that open-ended verbal alignment involves two complementary computations. First, context-dependent statistical prediction constrains the open lexical space to a set of plausible candidates. On each trial, this process was operationalized using a DLM (GPT-2), which generated a candidate word set for trial *t*+1 from the preceding conversational context and assigned prior selection probabilities to the candidates (Fig. 3b). Second, partner-directed pragmatic inference adjusts prior selection probabilities within this contextually constrained set according to the evolving interaction. Specifically, the model predicts the participant’s word-selection bias (WSB) for the next trial by updating their current bias in light of the partner’s responses (Fig. 3c). This predicted bias was used to reweight the DLM-derived prior probabilities, increasing the probabilities of candidates consistent with the predicted bias and decreasing those of competing alternatives (Fig. 3a). The resulting posterior distribution over candidate words defines the participant’s hypothesis landscape, in which the candidate set specifies the contextually relevant semantic search space and the posterior probabilities specify the relative selection priorities within that space. Through these complementary processes, the two participants were expected to coordinate their word choices within their respective hypothesis landscapes, progressively achieving precise lexical alignment.

**Fig. 3.**
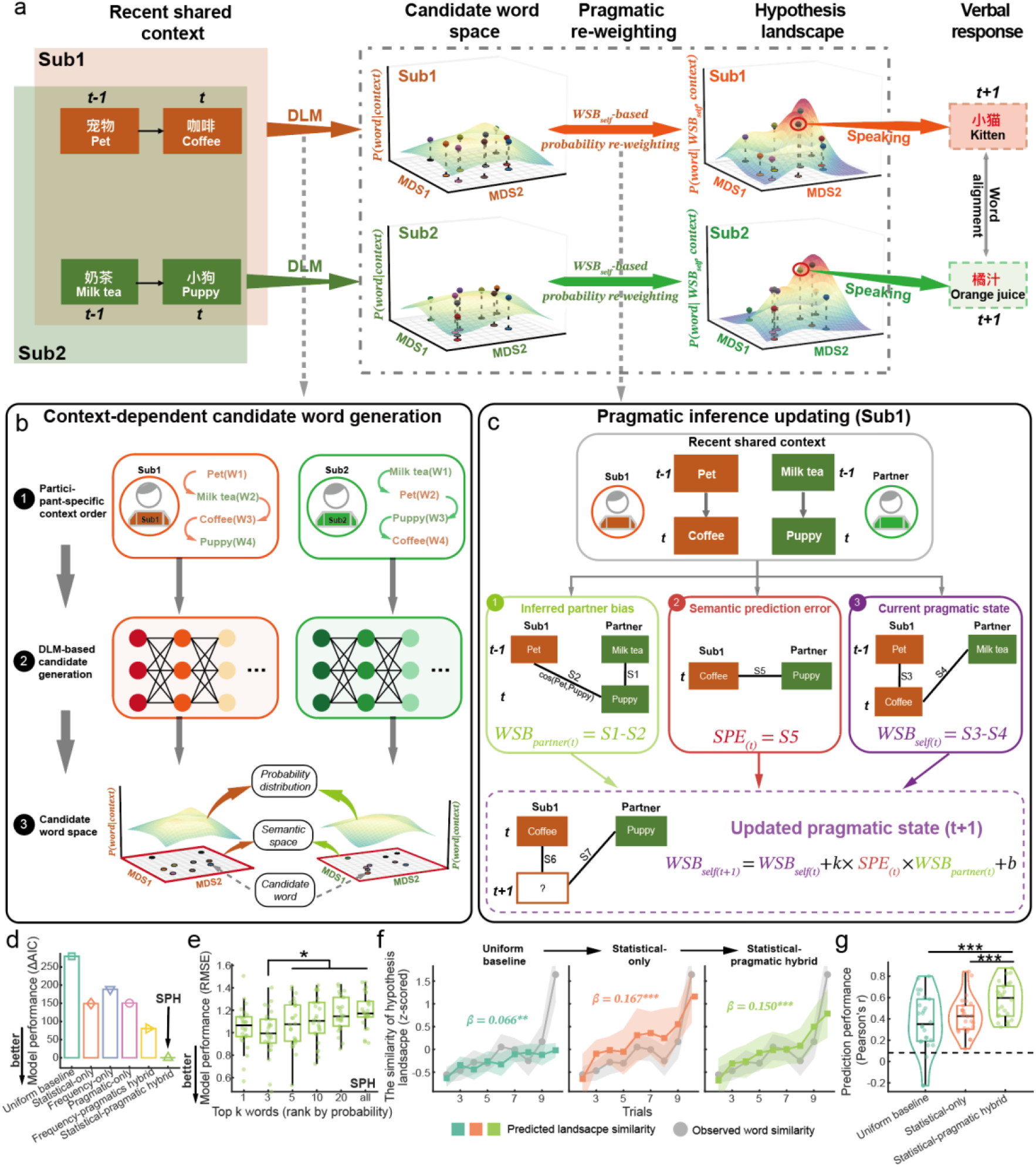
(a) Schematic of the word selection model. Word space comprising a two-dimensional semantic space, obtained via multidimensional scaling (MDS1 and MDS2), and an additional third dimension that represents the selection probability of each word. (b) The schematic of word space generation by DLM-based statistical prediction. (c) The structure of pragmatic prediction models from the perspective of subject 1. The full model space was empirically tested (seven models in total) but only the equation of the winning model is shown here (see Supplementary Text 4 for details). The model comprises three latent variables: inferred partner’s word selection bias (*WSB*_*partner(t)*_), semantic prediction error (*SPE*), and individual’s pragmatic prediction state (*WSB*_*Self(t)*_ and *WSB*_*Self(t+1)*_). (d) AIC-based model comparison in the *Exact communication* task. (e) The performance of the SPH model in the *Exact communication* task when different sizes of candidate clusters were included. (f) Trial-wise changes in observed semantic similarity and hypothesis-landscape similarity generated by the Uniform baseline, Statistical-only, and SPH models. Observed semantic similarity was identical across models. (g) Leave-one-dyad-out prediction of observed semantic similarity from the hypothesis landscapes generated by the Uniform baseline, Statistical-only, and SPH models, quantified as the Pearson correlation between predicted and observed values. The dashed line indicates the mean prediction performance across the three models under random shuffling. Abbreviations: Sub1/2: subject 1/2; W1-4: word1-4; ***w***: word; ***c***: context; DLM: deep language model. ^*^*p* < 0.05, ^***^*p* < 0.001, FDR corrected

Although the partner-sensitive component was specified a priori in SPH model, its updating rule was determined empirically. We formalized this component as a trial-by-trial updating process over word-selection bias, which captures the balance between maintaining one’s own preceding semantic trajectory and shifting toward the partner’s. Motivated by the AIF theory^24,25^, we compared seven candidate updating models that differed in whether and how the participant’s current selection bias was updated according to the inferred partner bias and semantic prediction error (SPE). Both in-sample and leave-one-dyad-out model comparisons identified the SPE-modulated partner-updating model (SPEMU) as the best-performing account in the *Exact communication* task (see Supplementary Text 4.3 for details). In this model, the inferred partner bias determined the direction of adjustment, whereas SPE modulated the magnitude of that adjustment. The cross-dyad parameters estimated from the remaining dyads were then used to generate out-of-sample pragmatic predictions for the held-out dyad, and this updating rule was incorporated as the pragmatic component of the SPH model.

#### Model comparison

We next asked whether lexical alignment was better explained by combining context-sensitive statistical prediction with partner-sensitive pragmatic reweighting than by either component alone or by simpler lexical alternatives. All models operated over the same DLM-generated candidate sets (see Fig. S5a–b and Supplementary Text 3 for details of the candidate sets and their lexical properties) and differed only in how probabilities were assigned to candidates. The Statistical-only model retained the DLM-derived contextual prior but omitted pragmatic reweighting, whereas the Pragmatic-only model assigned equal initial probabilities to candidates and relied only on partner-sensitive reweighting. Two additional models replaced the DLM-derived contextual prior with word-frequency-based lexical accessibility, either alone (Frequency-only) or combined with pragmatic reweighting (Frequency-pragmatics hybrid), and a Uniform baseline assigned equal probability to all candidates. Model comparison results in the *Exact communication* task showed that the SPH model performed best among all six models (Fig. 3d, lowest AIC, AIC weight SPH > 99.99%), indicating that error-driven pragmatic inference updating corrected the context-dependent statistical prediction prior during lexical alignment.

#### Landscape validation

Having established the relative advantage of the hybrid model, we next asked whether the resulting posterior distribution formed a behaviorally meaningful hypothesis landscape. We first tested whether candidates assigned higher posterior probabilities showed closer correspondence to participants’ actual lexical choices. Candidate words were ranked by their SPH-derived posterior probabilities to form top-k clusters (k = 1, 3, 5, 10, 20, or all). The embeddings within each cluster were averaged to obtain a semantic representation for each participant. For each dyad and each cluster size, model performance was assessed using the root mean squared error (RMSE, lower is better) across trials between the semantic similarity of the two participants’ semantic representations and that of their observed words. A one-way repeated-measures ANOVA revealed a significant main effect of cluster size (*F*(5, 25) = 7.987, *p* < 0.001, 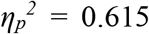, Fig. 3e). Further post-hoc comparisons revealed that the SPH model performed best when the top-3 words were included in the cluster, with top-3 performance significantly better than that of the top-5 to all-word sets (statistical details see Table S7), suggesting that the SPH posterior formed an ordered rank structure over the candidate sets.

Second, we tested whether the SPH-derived landscape became progressively aligned between interlocutors and whether landscape alignment predicted the observed lexical semantic alignment within the dyad. To this end, we calculated the trial-wise hypothesis landscape similarity within each dyad (see Methods). LME analysis revealed that the similarity of the SPH-based hypothesis landscape within dyads increased significantly across trials (*β* = 0.150, *t*(642) = 6.663, *p* < 0.001, *d* = 1.31, Fig. 3f). Moreover, cross-dyad linear regressions trained on the similarity of the SPH-based hypothesis landscape significantly predicted semantic similarity between the observed words, with an average *r* = 0.572 (*p* < 0.001, permutation-based, FDR corrected, Fig. 3g). To determine whether this landscape–behavior correspondence could be explained by candidate-set overlap or statistical prediction alone, we repeated the same analyses using the Uniform baseline and Statistical-only models. Although both control models also showed increases in within-dyad landscape similarity across trials (*β*_*Uniform*_ = 0.066, *β*_*Statistical*_ = 0.167, both *p* < 0.05, Fig. 3f), their prediction of observed semantic similarity was significantly weaker. A repeated-measure ANOVA and pairwise comparison further confirmed that the SPH model’s landscape predicted the observed semantic similarity significantly better than both the Uniform baseline (Δ*r* = 0.211, *p* < 0.001, FDR corrected) and Statistical-only models (Δ*r* = 0.145, *p* < 0.001; FDR corrected; Fig. 3g) (Statistical details see Table S8).

Together, these results suggest that the SPH model captured a structured and behaviorally meaningful hypothesis landscape. Its internal probability ordering corresponded to participants’ lexical choices, and its interpersonal similarity predicted observed lexical alignment beyond candidate-set overlap or statistical prediction alone.

### Interaction reorganizes the hypothesis landscape across global and local scales

Having established the behavioral validity of the inferred landscape, we next asked how interaction reorganized its internal structure across complementary global and local scales (Fig. 4a). We distinguished the global lexical-semantic search region defined by the candidate set from the local selection structure defined by the partner-adjusted probability distribution over candidates. We then characterized how these two levels changed both within individuals and between interlocutors over the course of interaction.

**Fig. 4.**
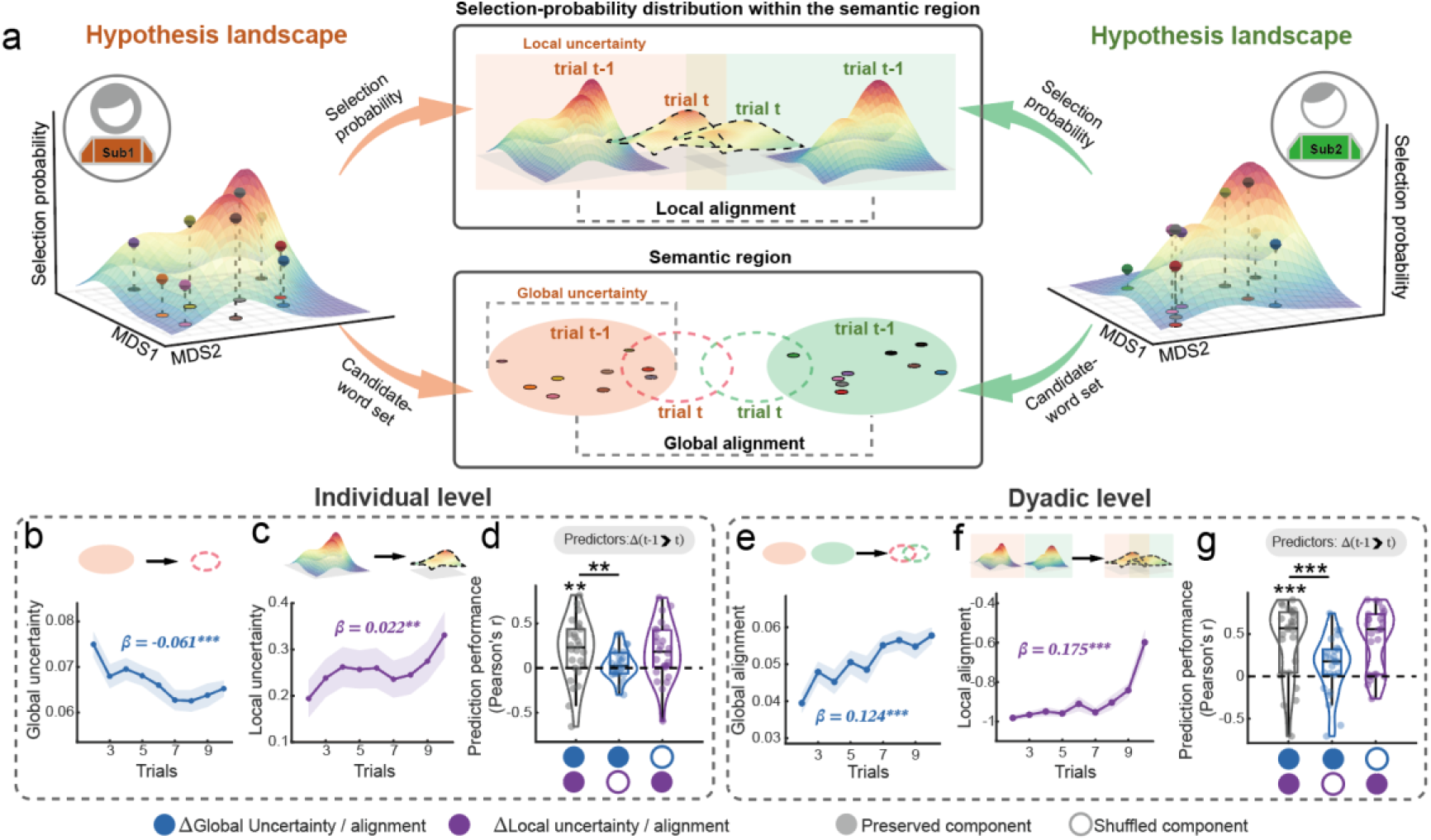
(a) Schematic representation of the hypothesis landscape for interacting interlocutors. For each interlocutor, candidate words are represented as points positioned according to their semantic relationships in a two-dimensional MDS space, with their overall spatial distribution defining the global lexical-semantic search region constrained by the preceding context. The height of the probability surface reflects the pragmatically adjusted selection probability assigned to each candidate word, with the distribution of these probabilities capturing local partner-sensitive selection priorities. (b–d) Individual-level landscape decomposition. Panels (b) and (c) show global uncertainty, reflecting the breadth of the candidate lexical-semantic search region, and local uncertainty, reflecting how distributed the pragmatically adjusted selection probabilities are across candidate words, respectively, across trials in the Exact communication task. (d) Leave-one-dyad-out prediction of subsequent lexical alignment from trial-to-trial changes in global and local uncertainty. (e–g) Dyadic-level landscape decomposition. Panels (e) and (f) show global alignment, reflecting the similarity between interlocutors’ candidate lexical-semantic search regions, and local alignment, reflecting the similarity between their pragmatically adjusted selection probability distributions, respectively, across trials in the Exact communication task. (g) Leave-one-dyad-out prediction of subsequent lexical alignment from trial-to-trial changes in global and local alignment. In (d) and (g), filled and open circles indicate preserved and shuffled predictors, respectively, and the dashed line denotes prediction performance when both predictors were shuffled. ^**^*p* < 0.01, ^***^*p* < 0.001, FDR corrected.

At the individual level, we first examined the complementary dynamics of global and local uncertainty. Global uncertainty captured the breadth of a participant’s lexical-semantic search regions and was quantified as the average semantic distance (1-cosine similarity) among all pairs of candidate words within a participant’s candidate set. Local uncertainty captured how distributed the pragmatically adjusted selection probabilities were across candidate words and was quantified as the normalized entropy of the selection probability distribution (normalized by the logarithm of the number of candidate words). Higher global uncertainty indicates a broader lexical-semantic search region, whereas higher local uncertainty indicates that selection remains distributed across multiple plausible candidate words rather than concentrated on a single dominant option. LMEs revealed that global uncertainty decreased across trials (*β* = −0.061, *t*(422) = −3.651, *p* < 0.001, *d* = −0.506, Fig.4b), indicating progressive narrowing of the lexical-semantic search regions during interaction. In contrast, SPH-derived local uncertainty increased across trials (*β* = 0.022, *t*(421) = 2.962, *p* = 0.003, *d* = 0.411, Fig. 4c). Moreover, although the Statistical-only model shared the same candidate word sets with the SPH model, its local uncertainty remains stable across interaction (*β* = 0.004, *t*(421) = 0.988, *p* = 0.324, *d* = 0.137; Fig. S7). Thus, interaction did not simply reduce uncertainty uniformly but redistributed uncertainty across scales: the broad semantic search region progressively narrowed, while selection remained distributed across multiple locally plausible alternatives.

We next asked whether dynamic fluctuation in global and local uncertainty influences subsequent lexical alignment in the *Exact communication* task. To this end, for each dyad, we quantified trial-to-trial changes in global and local uncertainty from trial (t-1) to trial (t) and used these changes to predict subsequent lexical alignment under leave-one-dyad-out cross-validation (Fig. 4d). The results showed that the full model, including both factors, significantly predicted subsequent lexical alignment, outperforming the all-shuffled baseline (Δ*r* = 0.198, *p* = 0.006). Critically, the full model significantly outperformed the local-uncertainty-shuffled model (Δ*r* = 0.150, *p* = 0.007), but not the global-uncertainty-shuffled model (Δ *r* = 0.016, *p* = 0.305). Together, these results indicate that subsequent lexical alignment was predicted primarily by prior changes in local uncertainty.

At the dyad level, we tested whether interlocutors became increasingly aligned in both the broader lexical-semantic regions they searched and the local selection priorities they assigned within those regions. Global alignment was quantified as the semantic similarity (cosine similarity) between the participants’ candidate word sets, whereas local alignment was quantified as the negative Jensen–Shannon divergence between their selection probability distributions within the shared candidate set (see Methods). LMEs revealed that both global and local alignment increased over trials in the *Exact communication* task (global alignment: *β* = 0.124, *t*(198) = 5.666, *p* < 0.001, *d* = 1.11; local alignment: *β* = 0.175, *t*(198) = 7.825, *p* < 0.001, *d* = 1.53; Fig. 4e-f). These results suggest progressive alignment at both the level of global lexical-semantic search regions and local partner-sensitive selection priorities.

The simultaneous increase in global and local alignment raised a further question: which level of interpersonal convergence was most consequential for what the dyad said next? To address this, we used trial-to-trial changes in global and local alignment to predict subsequent lexical alignment under leave-one-dyad-out cross-validation (Fig. 4g). In the *Exact communication* task, a full model including both factors significantly predicted the observed lexical alignment under leave-one-dyad-out cross-validation, outperforming the all-shuffled baseline (Δ*r* = 0.428, *p* < 0.001). Additionally, the full model significantly outperformed the local-alignment-shuffled model (Δ*r* = 0.293, *p* < 0.001), but not the global-alignment-shuffled model (Δ*r* = −0.030, *p* = 0.792). Thus, lexical alignment in *Exact communication* was predicted primarily by local alignment in partner-adjusted selection probabilities.

Together, these findings suggest a division of labor between the two levels of interpersonal coordination. Shared conversational context progressively brought interlocutors into increasingly similar lexical-semantic regions, but precise lexical convergence depended most strongly on their becoming aligned in how they prioritized alternatives within those regions. In this sense, global alignment appears to establish a shared search space, whereas local alignment provides the more proximal computational basis for selecting the same lexical outcome.

### Task specificity of global–local landscape organization

We next asked whether the global–local organization identified above reflected a general consequence of repeated lexical production or was selectively enhanced when interlocutors had to achieve precise lexical coordination. We fitted the Uniform baseline, Statistical-only and SPH models separately in each task. We first quantified the contribution of context-dependent statistical prediction as the performance difference between the Statistical-only and Uniform baseline models (indexed by RMSE). A one-way repeated-measures ANOVA revealed no significant main effect of task (*F*(2, 25) = 2.681, *p* = 0.088, 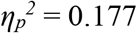, Fig. 5a), suggesting that the contribution of statistical prediction did not differ reliably across tasks. We then quantified the corrective contribution of error-driven pragmatic inference as the performance difference between the SPH and Statistical-only models. In contrast, this analysis revealed a significant main effect of task (*F*(2, 25) = 7.151, *p* = 0.003, 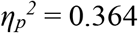, Fig. 5b). Further post-hoc comparisons showed that the active-inference corrective effect was significantly stronger (more negative) in the *Exact communication* task than in either control task (*Exact vs Category: t*(25) = −3.625, *p* = 0.004, *d* = −0.530; *Exact vs Solo: t*(25) = −2.960, *p* = 0.010, *d* = −0.701). No significant difference was found between the two control tasks (*p* > 0.05). These results suggest that DLM-based statistical prediction contributed similarly across tasks, whereas the corrective contribution of pragmatic inference was selectively enhanced during *Exact communication*.

**Fig. 5.**
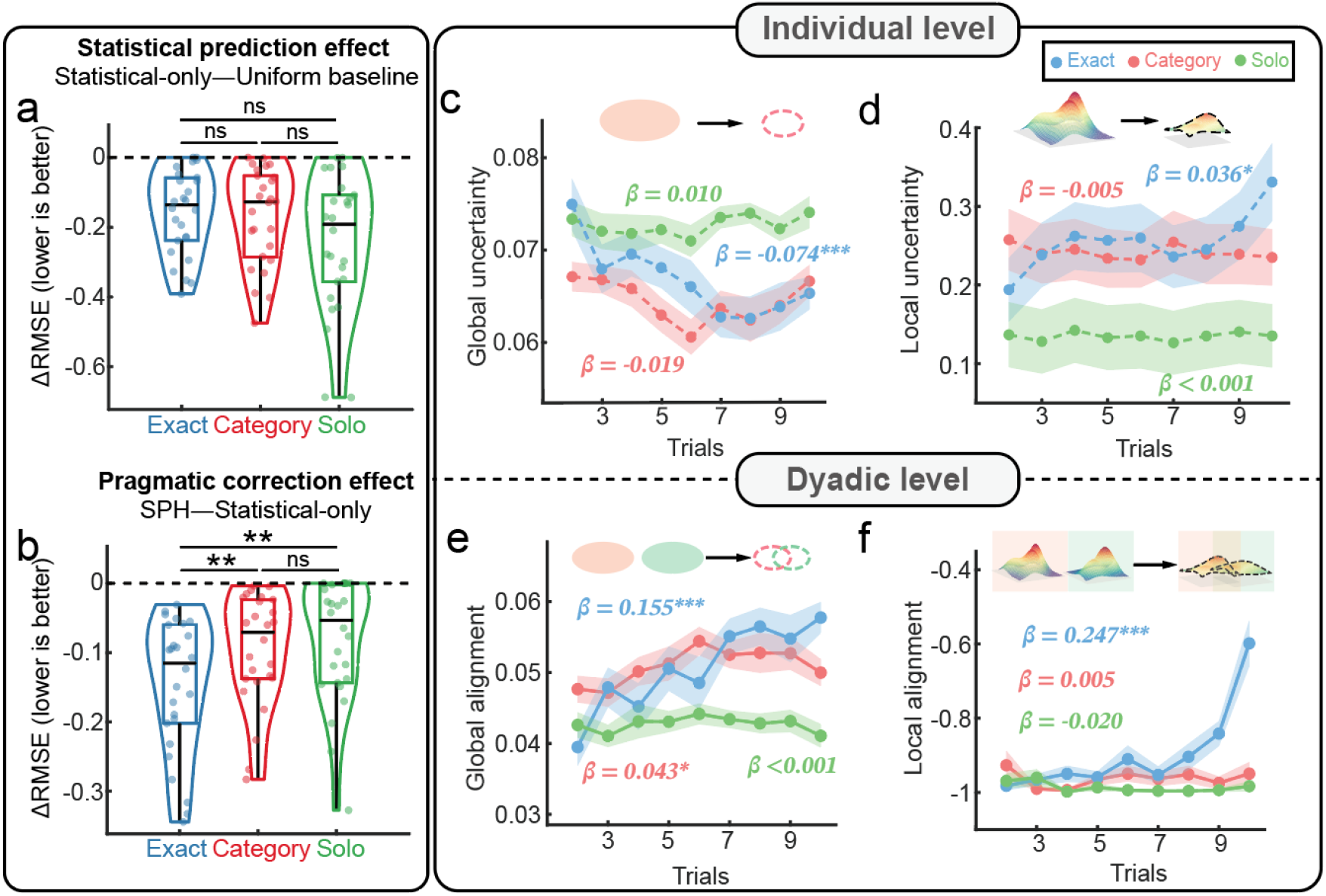
(a) Contribution of context-dependent statistical prediction across tasks, quantified as the difference in root mean squared error (RMSE) between the Statistical-only and Uniform baseline models. (b) Contribution of pragmatic correction across tasks, quantified as the RMSE difference between the SPH and Statistical-only models. (c) Global uncertainty across trials and tasks, reflecting the breadth of the candidate lexical-semantic search region. (d) Local uncertainty across trials and tasks, reflecting how distributed the partner-adjusted selection probabilities were across candidate words. (e) Global alignment across trials and tasks, reflecting convergence between interlocutors’ lexical-semantic search regions. (f) Local alignment across trials and tasks, reflecting convergence between interlocutors’ partner-adjusted selection probabilities. In all box plots, the center line indicates the median and the whiskers extend to the most extreme values within 1.5 times the interquartile range. The top and bottom of the boxes correspond to the third and first quartiles, respectively. ^*^*p* < 0.05, ^**^*p* < 0.01, ^***^*p* < 0.001, all two-sided, FDR corrected. SPH: statistical-pragmatic hybrid.

Next, we asked whether the global and local uncertainty dynamics identified in the previous section were also specific to *Exact communication*. LME analysis of global uncertainty revealed a significant trial-by-task interaction (*F*(2,1353.2) = 6.634, *p* = 0.001, 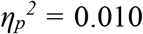, Fig. 5c), with simple slope tests showing a decrease across trials only in the *Exact communication* task and no significant change in the control tasks (see Table S9 for statistical details). For local uncertainty, the trial-by-task interaction was not significant (*F*(2,1353.1) = 1.986, *p* = 0.138, 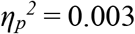, Fig. 5d). Simple-slope analyses indicated a significantly gradual increase across trials only in the *Exact communication* task (see Table S9 for statistical details). Together, the complementary dynamics between global and local uncertainty were specific to precise lexical communication.

Moreover, to determine whether changes in global and local uncertainty predicted subsequent lexical alignment in the control tasks, we repeated the leave-one-dyad-out prediction analysis using prior changes in global and local uncertainty as predictors (Fig. S8). Unlike in *Exact communication*, the full model, including both predictors, failed to outperform the all-shuffled baseline or either of the selective-shuffling models (all *ps* > 0.05). Thus, the predictive relationship between prior changes in local uncertainty and subsequent lexical alignment was evident in *Exact communication* but not in the control tasks.

We next examined whether global and local alignment of the SPH-derived dyadic hypothesis landscape was likewise modulated by task. To test this, we employed LMEs to examine trial-wise changes in the corresponding global and local alignments across tasks. Both global and local alignment showed a significant trial-by-task interaction (global alignment: *F*(2, 661.10) = 13.530, *p* < 0.001, 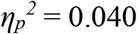; local alignment: *F*(2, 661.76) = 41.827, *p* < 0.001, 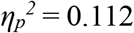), with stronger trial-wise increases in the *Exact communication* task than in the control tasks (Fig. 5e-f; see Table S10 for statistical details). These results indicate that *Exact communication* promoted progressive alignment in both the global lexical-semantic search level and the local partner-sensitive selection level.

Finally, we asked whether these two dyadic alignment measures contributed differently to the prediction of observed lexical alignment in control tasks (Fig. S8). The full model, including both predictors, significantly predicted subsequent lexical alignment in both tasks (*Category*: Δ*r* = 0.339, *p* = 0.001; *Solo*: Δ*r* = 0.260, *p* = 0.001, relative to all-shuffled baseline). Importantly, the full model significantly outperformed the global-alignment-shuffled model (*Category*: Δ*r* = 0.358, *p* < 0.001; *Solo*: Δ*r* = 0.258, *p* < 0.001), but not the local-alignment-shuffled model (all *ps* > 0.05). Thus, unlike in *Exact communication*, subsequent lexical alignment in the control tasks was predicted primarily by prior changes in global alignment rather than local alignment.

Together, these findings suggest that the contribution of context-dependent statistical prediction generalized across both interactive and non-interactive settings, whereas the error-driven pragmatic correction effect was selectively enhanced when communication required precise alignment of communicative actions. Under this communicative demand, the hypothesis landscape showed stronger global–local reorganization, with subsequent lexical convergence primarily predicted by local partner-sensitive dynamics.

### Neural organization of partner-sensitive pragmatic updating

The preceding results showed that error-driven pragmatical adjustment contributed more strongly to lexical selection during *Exact communication* than in control tasks. We therefore asked whether these process-level computations showed corresponding neural signatures during *Exact communication*. To address this question, we first examined the neural correlates of three key process-level components: an inferred partner’s word selection bias (WSB_partner_), semantic prediction error (SPE), and the subsequent partner-directed updating of one’s own word-selection bias (ΔWSB_self_). These quantities correspond to components predicted by the active-inference-motivated updating scheme used in the SPH model. The general linear model (GLM) analysis revealed a task-selective neural dissociation: the left TPJ and posterior temporal cortex tracked the inferred partner’s word-selection bias, the left anterior temporal cortex (aTC) encoded semantic prediction error, and the left inferior frontal cortex (IFC) tracked the subsequent updating of the participant’s own word-selection bias (Fig. S9 a-c and Table S11-12, see Supplementary Text 6.1 for more details).

Having identified these component-related regions, we next examined their functional integration across the thinking and speaking phases. During *Exact communication*, activity in the partner-directed inference (left TPJ) and semantic-prediction-error regions (aTC) during the late thinking phase (~4s) jointly predicted subsequent activity in the left IFC, a region associated with updating of the participant’s own word-selection bias during speaking. This temporally specific relationship was absent in the two control tasks, suggesting that partner-related information and semantic prediction error are integrated to support the updating of lexical-selection bias during precise verbal collaboration (see Supplementary Text 6.2 and Fig. S9 d-f for details).

Finally, we used an independent hierarchical localizer task to examine whether these functionally related regions operated at different levels of linguistic processing. The results showed that the left IFC associated with pragmatic-prediction updating (ΔWSB_self_) and the left TPJ associated with inferring the partner’s action (WSB_partner_) responded more strongly to larger linguistic units, such as narratives and sentences, while the left aTC associated with SPE responded more strongly to smaller linguistic units, such as words (see Fig. S11b-c and Table S14-15). These findings are consistent with a hierarchical dissociation in which higher-level regions support partner inference and partner-directed updating, whereas lower-level temporal regions encode the resulting semantic prediction error (see Supplementary Text 6.3 for more details).

### Neural representation of the hypothesis landscape

In the SPH model, collaborative word generation during *Exact communication* was characterized by a task-specific hypothesis landscape. We therefore asked whether this model-derived hypothesis landscape had corresponding neural correlates. To address this question, we employed representational similarity analysis (RSA), which is well suited to testing whether the structure of a high-dimensional hypothesis landscape is mirrored in neural activity (Fig. 6a–c; see Methods for details). Specifically, for each channel, we compared the trial-by-trial brain representational dissimilarity matrix (RDM) with the RDM derived from the SPH-based hypothesis landscape. Channel-wise RSA values from the two control tasks were averaged as a baseline, and *Exact communication* was compared against this baseline using a permutation-based null distribution of correlation differences, with *p*-values corrected across channels. The results revealed a significantly larger RSA correlation for the participant’s own hypothesis landscape than baseline in the left inferior parietal–supramarginal region (left IPL/SMG) (CH7) (Δ*r* = 0.103, *p* = 0.044; Fig. 6d-e). No significant effects were observed for partner’s hypothesis landscape (all *ps* > 0.05; Fig. 6f-g).

**Fig. 6.**
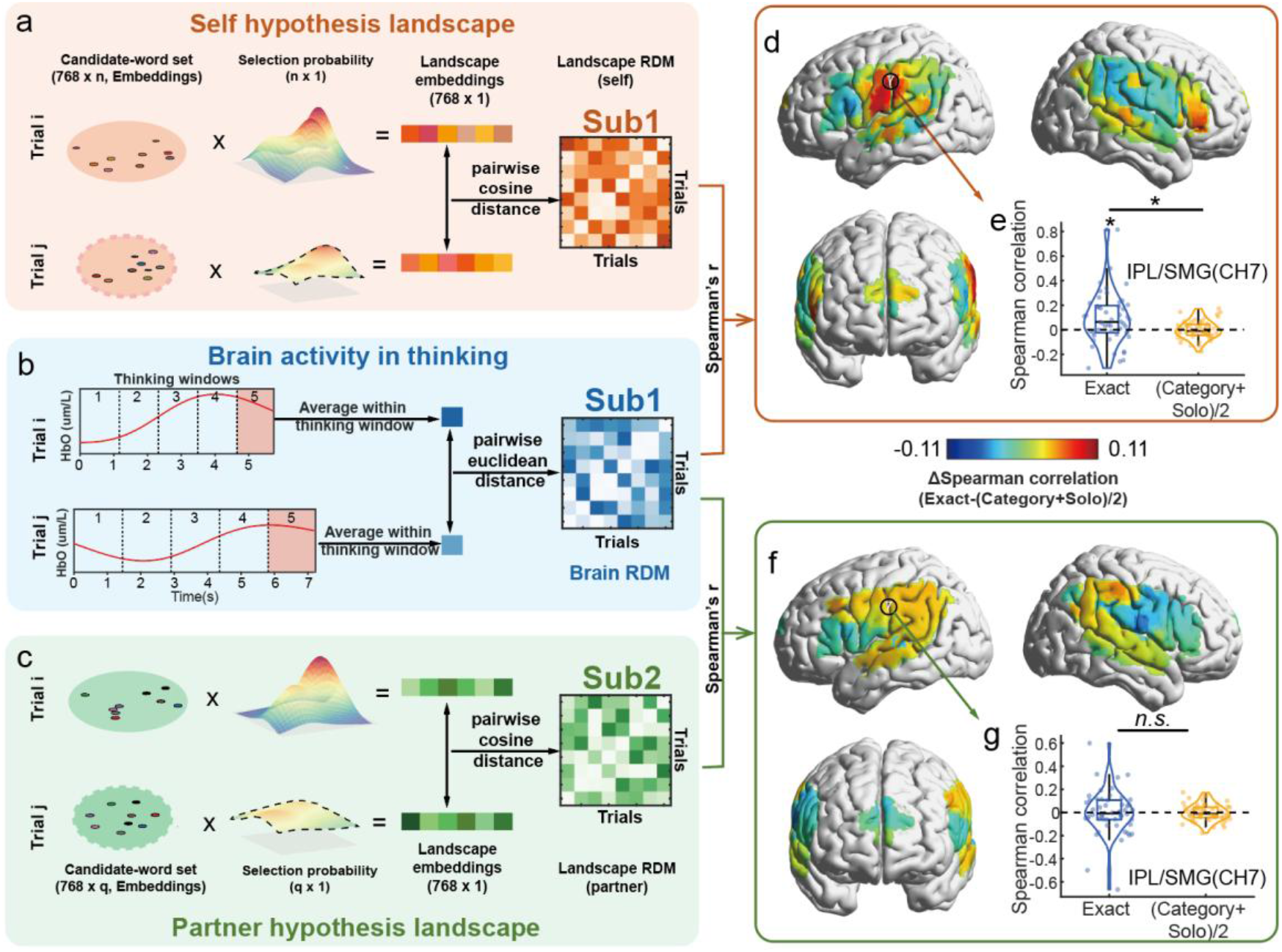
(a) and (c) The self- and partner-related landscape RDMs were constructed by computing the cosine distance between posterior-probability-weighted word embeddings across trials. (b) The brain RDM was constructed by computing the Euclidean distance between mean HbO responses within the fifth thinking window for each pair of trials. (d) and (f) Differences in Spearman RSA correlation coefficients between the *Exact Communication* and baseline tasks for the self-related (d) and partner-related (f) landscape RDMs. (e) and (g) RSA correlations between the landscape RDMs and the brain RDM at Channel 7, shown separately for the *Exact Communication* and baseline tasks for the self-related (e) and partner-related (g) landscapes. Sub, subjects; Win: windows; RDM: representational dissimilarity matrix; HbO: changes in oxyhemoglobin. ^*^*p* < 0.05, FDR-corrected; n.s., not significant.

To ensure that the observed task difference was not driven by negative RSA values in the control conditions, we further compared the channel-wise RSA values in the *Exact communication* task with a permutation-based null distribution (one-tailed, FDR-corrected). This analysis again revealed significant effects for one’s own hypothesized hypothesis landscape in the left IPL/SMG (CH7, *r* = 0.102, *p* = 0.022) and postcentral cortex (CH15, *r* = 0.101, *p* = 0.022; Fig. S12), whereas no significant effects were observed for the partner’s hypothesis landscape (*ps* > 0.05; Fig. S12).

Finally, to test whether these RSA effects could be explained by candidate word sets or by statistical prediction alone, we repeated the same analyses using the Uniform baseline and Statistical-only models. Neither model showed significant RSA correlations or between-task differences (*ps* > 0.05, FDR-corrected; Fig. S13). Together, these findings suggested that the SPH-derived hypothesis landscape was reflected in neural representational structure, with the most consistent evidence observed for the participant’s own landscape in the left IPL/SMG during *Exact communication*.

### Replication across model implementations and datasets

To assess the robustness and generalizability of the main findings, we conducted two complementary replication analyses. First, we asked whether the observed computational results depended specifically on GPT-2, which supports next-word prediction through relatively rich contextual semantic representations learned during pre-training. To address this question, we repeated the same analyses using the Chinese version of FastText, which provides semantic representations that are substantially less context-sensitive than those derived from GPT-2. The main findings were well replicated, confirming that the key computational patterns were robust to the choice of semantic representation (see Supplementary Text 7 for details).

Second, we asked whether the main findings generalized beyond the original sample and language context. To test this, we analyzed a publicly available independent dataset comprising 20 Czech-speaking dyads who engaged in an experimental task similar to the *Exact communication* task^43^. Because the Czech words produced by participants were translated into English, we applied the English version of GPT-2 to this dataset (see Methods). The same analytic procedures yielded convergent results, providing independent support for the main behavioral and computational findings on semantic convergence, hybrid-model superiority, and task-specific communicative-landscape organization (see Supplementary Text 8 for details)

Together, these replication analyses indicate that the core findings were robust across both model implementations and datasets. The contribution of active-inference-based correction did not depend on a specific DLM instantiation, and the main results generalized to an independent sample acquired in a different linguistic context.

## Discussion

Dynamic, open-ended communication requires interlocutors to select utterances that fit the immediate context while also adjusting to a partner whose intentions change across turns. To examine how this is achieved, we tested a hybrid computational division in which DLM-based statistical prediction generates a context-sensitive candidate word space, while partner-sensitive pragmatic inference adjusts selection priorities within that space through error-driven updating. Behavioral, computational, and neural findings were consistent with this hybrid computational account. In the following sections, we discuss these findings in turn.

First, previous studies have examined communicative convergence in relatively constrained communicative settings where utterance choices or descriptive targets are predefined^45,46^. Our results extend this line of work to open-ended communication, showing that progressive alignment can emerge even when the available information is sparse and the task provides no fixed alternatives in advance. Critically, this alignment was clearly task-dependent, suggesting that it reflected the demand for precise verbal collaboration rather than repeated word generation or broad semantic association.

Second, prior work has supported two important but complementary views of communicative alignment. One line of research suggests that human language processing exhibits DLM-like context-dependent predictive dynamics, whereby preceding linguistic context progressively constrains predictions about upcoming words and shapes their context-specific semantic representations^13,15,47^. Another has emphasized that successful communication requires pragmatic reasoning about a partner’s beliefs, intentions, and likely interpretations^7,21,48^. Recent neuro-symbolic approaches have begun to integrate these capacities within a hybrid computational framework, using language models to propose and evaluate open-ended candidate utterances or communicative hypotheses while rule-based cognitive models implement the pragmatic selection process^31,32^. Our findings suggest a different, process-level functional division. Context-dependent statistical prediction addresses the global search problem by constraining the otherwise open-ended utterance space to a context-relevant candidate region, whereas partner-directed pragmatic inference addresses the local selection problem by dynamically adjusting the relative priorities of candidates within that region across turns. Thus, statistical prediction and pragmatic inference may contribute to communicative alignment not simply by combining two sources of information, but by solving complementary computational problems at different scales: the former structures where to search, whereas the latter refines what to select for a specific partner.

Third, this computational division of labor was reflected in the internal dynamics of the hypothesis landscape. At the dyadic level, both global lexical-semantic search regions and local partner-sensitive selection priorities became increasingly aligned between interlocutors, with subsequent lexical convergence predicted primarily by local alignment. Consistent with active inference accounts of communication^24,25,49,50^, this pattern suggests that prediction-error-based pragmatic updating provides a mechanism for fine-grained partner-specific coordination, progressively aligning the relative priority of locally plausible alternatives, while shared contextual experience promotes convergence of broader lexical-semantic search regions.

Importantly, the individual-level dynamics revealed that global lexical-semantic compression was accompanied by increased local selection uncertainty, suggesting that the hypothesis landscape was reorganized rather than uniformly simplified during interaction. This pattern echoes previous findings in lexical production that selection becomes more competitive as semantically related alternatives become more closely clustered^37,38^. Our findings extend this principle to continuous collaborative communication, showing that increasing global constraint can coexist with increasing local competition and thereby revealing a dynamic balance between global semantic-search compression and local lexical competition. Moreover, the persistence of several locally viable candidates may reflect the availability of multiple lexical alternatives that can convey the same communicative intention^51^, thereby preserving expressive flexibility while allowing partner-specific inference to guide the final selection among these alternatives. Finally, this interpretation is further supported by the task-dependent pattern: the combination of progressive global compression and increasing local competition was evident when interlocutors were required to achieve exact lexical convergence, suggesting that the internal reorganization of the hypothesis landscape is sensitive to specific communicative goals.

Finally, the neural findings provided converging evidence for a distributed neural implementation of the hybrid statistical–pragmatic computations that support precise word selection during two-way open-ended communication. Activity in TPJ, anterior temporal cortex, and inferior frontal cortex tracked partner-related inference, semantic prediction error, and the updating of one’s own lexical-selection bias, respectively. This functional mapping accords with classic accounts linking TPJ and posterior temporal regions to mental-state reasoning and communicative perspective taking^52,53^, anterior temporal cortex to semantic representation^54,55^, and inferior frontal cortex to controlled semantic retrieval and selection among competing semantic representations^56,57^. Together, these correspondences suggest that precise open-ended communication may depend on both language systems that structure candidate meanings and social-inferential systems that track a partner’s communicative perspective.

Beyond this functional dissociation, an independent language localizer suggested a hierarchical organization of these component processes. Regions associated with partner inference (TPJ) and self-directed lexical-selection-bias updating (IFC) showed greater sensitivity to larger linguistic units, whereas the region associated with semantic prediction error (aTC) showed greater sensitivity to smaller, word-level units. Moreover, the selective RSA effect for the participant’s own hypothesis landscape in the left inferior parietal–supramarginal region suggests that the SPH-derived landscape was not merely a useful computational hypothesis, but was also reflected in neural representational structure. Previous work has implicated this region as an interface between lexical-semantic processing and perspective taking^58,59^. This finding suggests that the left IPL/SMG may link partner-sensitive pragmatic inference with lexical choice by representing a socially constrained, pragmatically weighted hypothesis landscape rather than a purely statistical semantic space.

It is important to acknowledge several limitations of the present study. First, the hybrid computational account was tested in a single-word communication task. As an initial step, this level of analysis provided a tractable way to study open-ended verbal coordination, while also offering a computational framework that may, in principle, extend to longer linguistic units, such as phrases, sentences, and discourse-level exchanges. Future work is needed to test this scalability directly by applying the present framework to more naturalistic communicative contexts involving extended linguistic sequences. Second, the construction of the candidate word space relied on DLM-based generation, and alternative approaches were not examined. Future work could extend the framework to alternative sources of lexical candidates, such as human word-association data. Third, although fNIRS provided a useful way to study spoken interaction under relatively natural conditions, its limited spatial resolution and penetration depth constrain anatomical precision and preclude direct observation of subcortical contributions. Future studies could extend the present findings using methods with higher spatial resolution, such as fMRI. As an initial step in this direction, we examined the generalizability of the main behavioral and computational findings in an independent dataset from a dual-fMRI study^43^. Building on this, it will be important to examine how subcortical regions, as well as cortical-subcortical interactions, contribute to the computational processes underlying partner-directed updating during communication.

In conclusion, the present study suggests that precise verbal collaboration in dynamic, open-ended communication depends on a hybrid computational process. Statistical prediction generates a context-sensitive candidate word space, whereas active-inference-inspired pragmatic adjustment refines lexical selection within that space through error-driven updating. This account helps explain how interlocutors can progressively achieve alignment under conditions in which relevant alternatives are not specified in advance and communicative demands change across interaction. By linking behavioral alignment, model-based evidence, and neural findings within a single framework, the present work provides a process-level account of how open-ended communication is organized in human interaction and may inform future studies of adaptive human–AI communication.

## Methods

### Participants

G^*^power 3.1 was used to determine the sample size of this study^60^. To achieve a power of 0.80 with a medium effect size (*Cohen f* = 0.25) and *α* = 0.05 in the major analysis of this study (repeated measure ANOVA), a minimum sample size of 24 dyads (48 participants) is required. To account for potential data loss, 30 dyads (60 healthy adults; 30 females; mean age = 20.5 years, SD ± 2.318) were finally recruited in Beijing through advertisements. These participants were randomly paired into 30 same-gender dyads. All participants were right-handed, had normal or corrected-to-normal vision, and had no history of neurological or psychiatric disorders. During data analysis, one participant missed over 10% of word recordings; consequently, the entire dyad’s data were excluded. Additionally, four participants’ brain data were removed due to the recording failure of the fNIRS system. As a result, each analysis included at least 52 participants.

The study protocol was approved by the Institutional Review Board of the State Key Laboratory of Cognitive Neuroscience and Learning, Beijing Normal University. Written informed consent was received from all participants.

### The collaborative word generation task

The collaborative word generation task (Fig. 1) was modified from a previous study^43^. Before the formal experiment, participants completed three practice rounds (five trials each) to familiarize themselves with the task procedure. Afterward, participants were required to verbally restate the task instructions to ensure comprehension. The formal experiment commenced only after the experimenter (T.Z.) verified that participants had accurately understood the task requirements.

The formal experiment included three tasks. In the *exact communication* task, two individuals of a dyad were seated in front of a computer in two separate rooms where they could not hear or see each other (Fig. 1a). Across 11 consecutive trials (1 round), they were requested to simultaneously produce the exact same Chinese words (1–4 characters), without repeating the word from the immediately preceding trial (Fig. 1c). In the first trial, participants had 5 s to think about what to say and another 3 s to produce the word aloud. The voices were recorded using a microphone (NESSIE USB Microphone, Blue Microphones Inc, USA, dual channel, 48k Hz) attached to a table. From the second trial onwards, at the start of the trial (before they began thinking of a new word), participants listened (3 s) to the audio of the voice recorded from their partner during the previous trial via an in-earphone (Xiaomi Corporation, China, dual channel). Subsequently, the duration of the thinking period was freely controlled by the participant and could be ended by pressing the key of “p” or “q”. Once both participants responded, they were required to articulate a new word within 3 s. There were no other restrictions on word generation, thereby creating an open-ended two-way verbal communication context. If the dyad reached precise semantic convergence before completing all 11 trials in a round, they were instructed to repeat the final word (identical for both members) for the remaining trials to match the trial count of other rounds. These repetition trials were not entered into later data analyses. The first trial was also excluded from later analysis due to the absence of prior conversational context.

Each dyad was additionally requested to perform two control tasks. In the *category communication* task, two individuals of a dyad were requested to say words that belonged to the same semantic category but not the same word as that of their partner. In the *solo production* task, the two members of a dyad were asked to independently say any words and the audio recording of the word produced by each individual was played back to them rather than their partner. The other part of the control tasks was the same as that in the *exact communication* task. Each task was repeated 4 times (4 rounds) with a 15-second resting-state interval between rounds that served as a baseline. The order of the three tasks was counterbalanced across dyads. The type of tasks and trial number were displayed in the top left corner of the screen to provide a prompt for participants. Two fNIRS systems were simultaneously employed to measure hemoglobin concentration changes in the brains of both dyad members during all three tasks (Fig. 1b).

### Coding of behavioral data

During the word generation task, words produced by participants were audio-recorded using a microphone. The epochs of the speaking and listening phases were extracted through manual annotation of the audio recordings using the Praat software (6.2.17). It should be noted that the period of listening in one member was the same as that of speaking in the other individual within a dyad in the *Exact communication* and *Category communication* tasks, or the same as speaking for the same individual during the *Solo production* task. Thus, only the durations of speaking were extracted; specifically, the period between the participant’s exhalation before speaking and the end of the sound waveform (Fig. S1). For the thinking phase, we marked the offset of the waveform from a preceding word as the onset and the time at which the participant pressed the key as the offset.

### Dynamic pattern of semantic convergence

To examine the probability of achieving precise semantic convergence in the *Exact communication* task, we counted the number of rounds in which participants met this task requirement. To further delineate the dynamic pattern of semantic convergence in each dyad across trials, we calculated the semantic similarity between words produced by two individuals of a dyad. Specifically, an Application Programming Interface (API) (https://cloud.baidu.com/product/speech) was employed to convert the audio of words into written texts. The transcription was manually checked and corrected by the experimenter (T. Z.). Subsequently, the transcribed texts were vectorized using the GPT-2 (Chinese version, https://huggingface.co/uer/gpt2-chinese-cluecorpussmall). In this process, because a single word might be segmented into multiple sub-word tokens by the model’s tokenizer, embeddings for all sub-word tokens were extracted and averaged to represent the semantic content of the word. Here, the ninth layer (twelve layers in total) was extracted as the embeddings of semantic features, in line with recent findings that intermediate layers of DLMs provide better semantic representations^61,62^. Based on word embeddings, semantic similarity was calculated trial-by-trial using cosine similarity, then minimal-centralized across tasks and dyads.

Additionally, since the number of trials in the *Exact communication* task varied across rounds due to exclusion of the first trial and trials involving word repetition, we resampled the trials in each round by performing percentile-based normalization to ensure that trials from rounds with fewer trials were proportionally mapped onto a standardized 10-trial scale. Rounds with fewer trials were padded with NaN. For example, if a dyad reached precise semantic convergence in 6 trials, the percentile of these trial positions was coded as [16.67%, 33.33%, 50%, 66.67%, 83.33%, 100%] (i.e. trial number/total number of trials before convergence^*^100%). Then, the first trial (at the 16.67th percentile) was mapped to the second position in the 10-bin sequence because it corresponded to the 10%–20% percentile bin. The same percentile-based alignment rule was applied to the remaining trials, resulting in a resampled sequence [NaN, trial 1, NaN, trial 2, trial 3, NaN, trial 4, NaN, trial 5, trial 6]. Finally, semantic similarity was averaged over 4 rounds of the same task. In the two control tasks, no resampling was performed; semantic similarity values in each trial were directly averaged across the 4 rounds. The LME analysis was conducted on the round-averaged semantic similarity, with *Task, Trial* and their interaction as fixed effects and *Dyad* identity as the random effect, while the durations of speaking and thinking were included as covariates.

### Candidate word-set generation

Motivated by prior work on the capacity of human working memory^63^ and the sequential nature of linguistic processing^64^, we constrained the DLM to use only the local conversational context while preserving the temporal order in which words were produced or heard by each interlocutor. Meanwhile, speaker identity (self vs. partner) was not encoded in order to minimize the influence of social information on this statistical prediction process. Specifically, for each trial, the four words from the preceding two trials were input into GPT-2 to generate the next candidate word set for that trial. For each participant within a dyad, the order of the preceding context words was consistent with the order in which the participant actually processed the conversational context. For instance, participant A’s context word order was [A_t-1_ B_t-1_ A_t_ B_t_], while that for participant B was [B_t-1_ A_t-1_ B_t_ A_t_]. The generated candidate words were required to have the same GPT-2 token length as the observed word on that trial, and the number of candidate words generated was set to 200 for each participant on each trial. Duplicate words, pseudowords (i.e. zero-frequency words), words repeated from the immediately preceding trial, and words containing punctuation were removed. Each remaining valid candidate word was retained with a transition score, which indexed its relative unnormalized support given the preceding context and was operationalized as the sum of the conditional logits assigned by GPT-2 to all tokens of the word across successive autoregressive transitions^12,65,66^. Additionally, given the open-ended nature of the collaborative word-generation task, participants had considerable flexibility in their lexical choices. Consequently, the 200 generated candidates may not always include the observed target words. For trials in which the target word was absent, we added it to the corresponding candidate set to ensure that this finite approximation of the lexical search space could accommodate participants’ observed choices (see Supplementary Text 3 for details).

To examine whether the words actually produced by human participants were consistent with candidate words generated by the DLM, linguistic attributes such as word frequency and POS were examined. Word frequency was defined as the occurrence frequency of candidate words in large-scale text corpora, while POS of words was defined as the words’ syntactic categories, both of which were obtained from the Natural Language Toolkit and “Jieba” packages in Python (https://github.com/fxsjy/jieba?tab=readme-ov-file). We conducted *χ*^2^ goodness-of-fit tests to determine whether the human-produced and DLM-generated words differed in their distributions of POS and word frequency within each task.

### Computational modeling of word selection

#### SPH model

The SPH model formulates trial-wise lexical selection during collaborative word generation as Bayesian inference over a dynamically generated candidate word set. The model predicts lexical selection on the upcoming trial t+1 using information available through trial t. For readability, the participant index is omitted below; all candidate sets, pragmatic states, and selection probabilities were computed separately for the two members of each dyad.

The model proceeds in three steps. First, the DLM uses the preceding conversational context to generate a candidate word set and assigns each candidate a context-dependent statistical prior. Second, partner-directed pragmatic inference updates the speaker’s pragmatic prediction state, and each candidate is evaluated according to how well it expresses this updated state. Third, the statistical prior and pragmatic likelihood are combined to yield the posterior probability of selecting each candidate.

##### Posterior probability

For candidate word 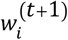 at trial *t* + 1, the posterior probability of selection was defined as

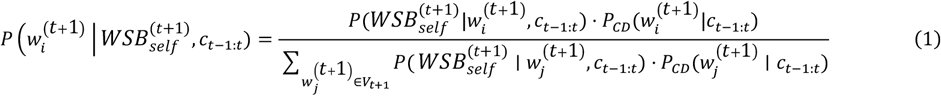

Here, 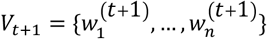 denotes the candidate word set at trial *t* + 1, *c*_*t*−1:*t*_ denotes the preceding four-word conversational context. 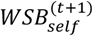 denotes the speaker’s updated pragmatic prediction state. The 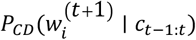 represents the DLM-derived context-dependent statistical prior, whereas 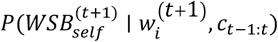, represents the pragmatic likelihood of candidate 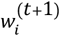. Thus, candidates receive higher posterior probabilities when they are both contextually plausible and compatible with the speaker’s updated pragmatic prediction state.

##### Context-dependent prior

The context-dependent prior 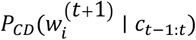 was derived from the transition score (logit) generated by the DLM for each candidate word given the preceding conversational context. These logits were transformed into normalized prior probabilities using a softmax function:

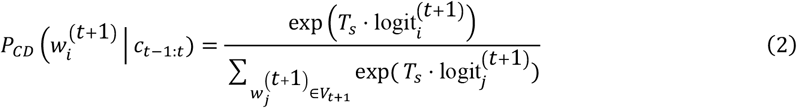

Here, *T*_*s*_ > 0 is a free temperature parameter controlling the sharpness of the prior distribution. Larger values of *T*_*s*_ produce a more concentrated distribution that assigns greater probability to candidates with higher DLM logits.

##### Pragmatic likelihood

The pragmatic likelihood 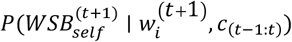 quantifies how well a candidate word 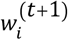 can convey the speaker’s updated pragmatic prediction state. To derive the pragmatic likelihood, we constructed a candidate-specific probability density over a continuous persistence–alignment dimension, which represents the balance between maintaining the speaker’s own preceding semantic direction and aligning with the partner. Each position along this dimension corresponds to a possible pragmatic state. The candidate-specific density was determined by two quantities: pragmatic utility (PU), which quantified the compatibility of a candidate with different possible pragmatic states, and pragmatic cost (PC), which reflected its confusability with competing candidates in the current candidate set (more distinctive words incur lower pragmatic cost). The speaker’s updated pragmatic prediction state 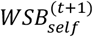 then specified a particular predicted state on this dimension for the upcoming trial, and the probability mass assigned by each candidate-specific density around this state determined its pragmatic likelihood.

1. For each candidate word 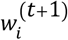, we then defined candidate-specific word-selection bias, 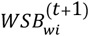, which located the candidate on the persistence–alignment dimension:

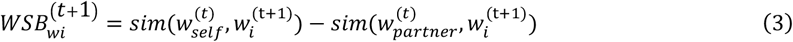

where *Sim*(.,.) denotes the semantic similarity (cosine similarity) between corresponding word embeddings. Here, 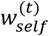 and 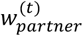 denote the words actually produced by the speaker and partner at trial *t*, respectively, whereas 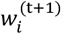 denotes a DLM-generated candidate word for the upcoming trial (Fig. 3c). Thus, 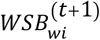 with more positive values were positioned toward self-persistence, whereas those with more negative values were positioned toward partner alignment.
2. Given its position on the persistence–alignment dimension, each candidate was evaluated in terms of PU and PC. PU measured the compatibility between the candidate-specific 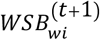 and a possible pragmatic state *x* on the persistence–alignment dimension:

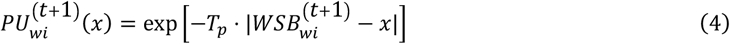

PC measured the confusability of the candidate with the other words in the same candidate set:

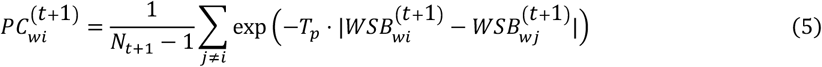

where *N*_*t*+1_ denotes the number of candidates in the candidate set and *T*_*p*_ > 0 is a free temperature parameter controlling sensitivity to pragmatic distance. Because the persistence–alignment dimension is one-dimensional, pragmatic distance was quantified as the absolute difference between two WSB values. Here, *x* denotes a possible pragmatic state along the continuous persistence–alignment dimension, rather than the speaker’s updated pragmatic prediction state on a particular trial.
3. PU and PC were then combined and normalized over the continuous range [*l, u*] of pragmatic-state dimension, yielding a candidate-specific probability density over possible pragmatic states:

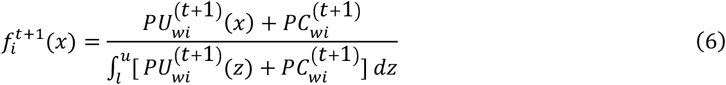

This density characterizes how strongly a given candidate supports different positions along the persistence–alignment dimension. Here, *l* and *u* were defined as the lower and upper bounds, respectively, of each individual’s word-selection bias values observed within each task. The denominator normalizes the density over [*l, u*] and admits a closed-form solution, which was evaluated analytically (see Supplementary Text 5). Because PC was constant with respect to *x*, larger PC values flattened the normalized density, making more confusable candidates less specific to any particular pragmatic state.
4. The speaker’s updated pragmatic prediction state, 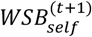, specified the particular predicted position on the persistence–alignment dimension for the upcoming trial. According to the selected pragmatic prediction model,

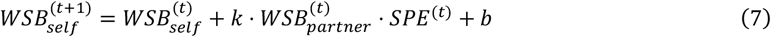

Here, 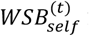 is the speaker’s previous pragmatic prediction state on the preceding trial, 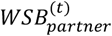 is the word selection bias derived from the partner’s preceding behavior, and *SPE*^(t)^ is the semantic prediction error at trial *t* (see definition in Supplementary Text 4.1, SPEMU model). The structure of this update equation was selected from seven candidate models. For each held-out dyad, the parameters *k* and *b* were estimated using data from the remaining dyads and were subsequently fixed before fitting the SPH lexical-selection parameters (Supplementary Text 4.3 for more details). The pragmatic likelihood of each candidate was then approximated by the local probability mass around the speaker’s updated pragmatic prediction state. Specifically,

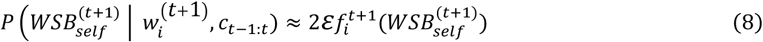

with *ε* = 0.005. Thus, candidates whose pragmatic-state densities assigned greater local probability mass around the speaker’s updated pragmatic prediction state received higher pragmatic likelihoods. The half-width *ε* was fixed across candidates and was not a free parameter. Because the resulting interval width 2*ε* was common to all candidates, this term cancels when the posterior is normalized across candidate words in Eq. 1 and therefore does not affect their relative posterior selection probabilities.

Finally, the posterior probability (Eq. 1) of lexical selection was obtained by normalizing the product of the context-dependent prior and the pragmatic likelihood across all candidate words.

##### Dyadic-level collaborative evaluation

Model fitting was performed at the dyadic level because the phenomenon of interest was communicative alignment between participants. We compared model-derived dyadic semantic similarity with the observed similarity between the two produced target words. For each participant and trial, we first summarized the model-predicted semantic representation as a posterior-weighted combination of the candidate-word embeddings. Let 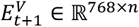 denote the embedding matrix of the candidate word set, and *p*_*t*+1_ ∈ ℝ^*n*×1^ denote the posterior probability vector from Eq. 1. The weighted embedding *h*_*t*+1_ was computed as

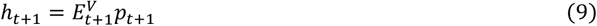

For each dyad, model-derived dyadic semantic similarity was quantified as the cosine similarity between the two participants’ posterior-weighted semantic representations *h*_*t*+1_:

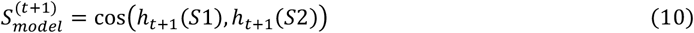

where *S*1 and *S*2 denote the two members of the dyad. This model-derived dyadic semantic similarity was compared with the observed semantic similarity between the words actually produced by the two participants,

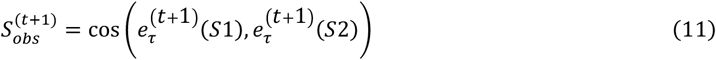

where 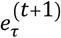 denotes the embedding of the target word actually produced at trial *t* + 1. Model parameters were estimated by minimizing the mean squared error between predicted 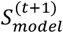 and observed dyadic semantic similarities 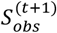:

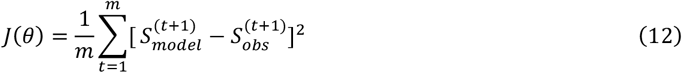

where *m* is the number of valid trials for each dyad and *θ* denotes the two free temperature parameters, *T*_*s*_ for statistical-prior sharpness, and *T*_*p*_ for pragmatic-distance sensitivity, estimated separately for each dyad.

#### Alternative models

To test whether lexical selection required the integration of context-dependent statistical prediction and partner-directed pragmatic inference, we compared the SPH model with five alternative models. All six models used the same participant- and trial-specific candidate word sets generated by the DLM, as well as the same dyadic-level fitting objective and optimization procedure. They differed only in the prior probabilities assigned to the candidates and whether these probabilities were subsequently adjusted by the pragmatic likelihood.

The Statistical-only and Pragmatic-only models tested whether either component of the SPH model was sufficient on its own. The Frequency-only and Frequency– Pragmatics Hybrid models tested whether the context-dependent DLM prior could be replaced by context-independent word frequency. The Uniform baseline model assigned equal probability to all candidates. Because all models used the same DLM-generated candidate sets, these comparisons concerned the assignment of probabilities within a candidate set rather than the generation of the candidate set itself.

#### Statistical-only model

The Statistical-only model tested whether lexical selection could be explained by context-dependent statistical prediction alone. Because no pragmatic reweighting was applied, the selection probability of each candidate was given directly by the DLM-derived prior:

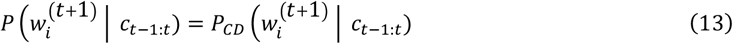

where 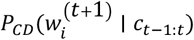was defined in Eq. 2 of the SPH model. The only free parameter was *T*_*s*_, controlling the sharpness of the DLM-derived prior, as defined in Eq. 2.

#### Pragmatic-only model

This model tested whether partner-directed pragmatic inference could guide lexical selection in the absence of graded statistical prior probabilities. It retained the pragmatic likelihood of the SPH model but replaced the context-dependent prior with a uniform prior over the DLM-generated candidate set:

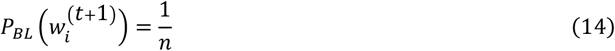

Where *n* is the number of candidate words in the set. Substituting this uniform prior into Eq. 1 meant that selection probabilities within the candidate set were determined entirely by the pragmatic likelihood. The only free parameter was *T*_*p*_, controlling sensitivity to pragmatic distance defined in Eqs. 4–5.

#### Frequency-only model

The Frequency-only model tested whether candidate selection could be explained by frequency-based lexical accessibility. The frequency-based prior was defined as:

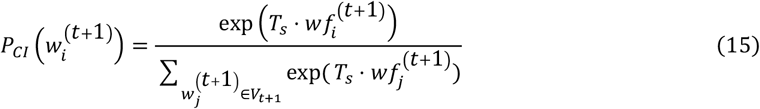

Here, 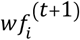 denotes the frequency score of candidate word *i*, and *T*_*s*_ > 0 is a free temperature parameter controlling the sharpness of the frequency-based prior. Because this model did not include pragmatic reweighting, its final selection probabilities were given directly by 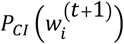. The only free parameter was *T*_*S*_, controlling the sharpness of the frequency-based prior.

#### Frequency-Pragmatics Hybrid model

This model retained the pragmatic likelihood of the SPH model but replaced the context-dependent DLM prior with a context-independent frequency-based prior defined in Eq.15. Specifically, 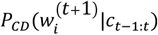 in Eq. 1 was replaced 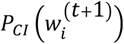 in Eq. 15 while all pragmatic terms remained unchanged. This model tested whether partner-directed pragmatic updating could improve lexical selection when the initial preferences among candidates reflected general lexical accessibility rather than the preceding conversational context.

The model contained two free parameters *T*_*s*_, controlling the sharpness of the frequency-based prior, and *T*_*p*_, controlling sensitivity to pragmatic distance. Although the Frequency-only and Frequency–Pragmatics Hybrid models used the same functional form for 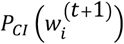, their parameters were estimated separately.

#### Uniform baseline model

The Uniform baseline model assigned equal selection probability to all words in the DLM-generated candidate set. It incorporated neither graded prior probabilities nor pragmatic reweighting and contained no free parameters.

#### Model fitting and comparison

All models were fitted to the behavioral data (i.e. word embeddings) using MATLAB 2020b with a custom function. The optimal parameters were estimated by minimizing *J*(*θ*) in Eq.12 (“fminsearch” function). Bounded free parameters were transformed from the [−∞: +∞] to the native model space via appropriate link functions (i.e. an exponential function was used to estimate the temperature parameters *T* to ensure positivity). In addition, to avoid the estimation being stuck in local minima, 50 different initial values were randomly set for the free parameters, from which the optimal parameters were selected.

For model comparison, first, AIC, and AIC weights were calculated as the index of model performance in the *Exact communication* task^67^. Lower AIC values and higher AIC weights indicate stronger relative support for a given model. Second, to compare the performance of the Statistical-only and SPH models across tasks, we fitted the Uniform baseline, Statistical-only, and SPH models to each control task. For each task, we subtracted the Uniform baseline model’s RMSE from that of the Statistical-only model to isolate the pure contribution of the statistical prediction mechanism in each task. A one-way repeated-measures ANOVA was then conducted to compare these performance differences across tasks, with speaking and thinking durations included as covariates. Analogously, we subtracted the Statistical-only model’s RMSE from that of the SPH model in each task to quantify the corrective effect of the active inference mechanism on the statistical prediction prior. A second one-way repeated-measures ANOVA was performed on these differences across tasks, again controlling for speaking and thinking durations as covariates.

#### The top-k word cluster analysis for model comparison and evaluation

To further investigate the word selection performance of each model, we examined whether words with higher selection probabilities were more similar to the target words than those with lower probabilities for each model. This analysis was conducted specifically in the *Exact communication* task. Candidates were ranked by model-derived selection probability, and top-*k* clusters were constructed for *k* = 1, 3, 5, 10, 20, and all candidates. For each cluster, candidate embeddings were averaged, and model performance was quantified as the RMSE between cluster-based and target-word semantic similarity for each dyad. Performance across cluster sizes was compared using a one-way repeated-measures ANOVA.

#### The hypothesis landscape analysis

To examine the trial-wise evolution of the hypothesis landscape within dyads, we analyzed the trial-wise dynamics of the model-derived landscape similarity, denoted here as landscape similarity (LS). For each participant, the hypothesis landscape was summarized by a posterior-probability-weighted embedding of the candidate word set. Specifically, 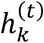 was computed as the weighted sum of the embeddings of all candidate words generated by the model for participant *k* at trial *t*, with weights given by their posterior lexical-selection probabilities. Thus, candidate words with higher posterior probability contributed more strongly to the final landscape representation (Eq. 9). Landscape similarity within each dyad was then defined as

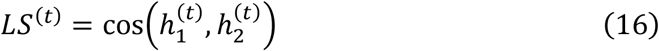

Higher LS values indicate greater convergence between the two participants’ internal hypothesis landscapes.

To examine the temporal dynamics of landscape similarity, *LS* was first computed at the trial level and then aligned using the same percentile-based normalization procedure as in the semantic similarity analysis. The aligned values were averaged across the four rounds of the same task for each dyad. LMEs were then fitted with aligned trial ID as a fixed effect and dyad identity as random effect. For model validation, the same procedure was repeated for the Uniform baseline, Statistical-only, and SPH models.

To test whether model-derived landscape similarity could explain the observed semantic similarity between the words produced by the two members of each dyad, we performed cross-dyad L1-regularized linear regression. Specifically, for each held-out dyad, the linear regression model was trained on the trial-wise data from all remaining dyads, using model-derived landscape similarity as the predictor and observed semantic similarity as the outcome. The trained model was then applied to the trial-wise landscape similarity of the held-out dyad to predict its observed semantic similarity trajectory. Prediction performance was quantified as the Pearson correlation between predicted and observed semantic similarity.

#### Global and local dynamics within the hypothesis landscape

To characterize the internal dynamics of the hypothesis landscape generated by the SPH model, we quantified trial-by-trial changes in collaboration and uncertainty at two levels: the overall structure of the candidate set, and the distribution of lexical selection within that set. For each participant *k* ∈ {1,2} and trial *t*, the SPH model generated a candidate word set 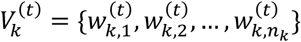 and a posterior word-selection probability distribution 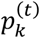 over that set. Let 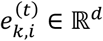 denote the semantic embedding of the *i*-th candidate word for participant *k* at trial *t*, where *n*_*k*_ is the number of candidate words. All candidate-word embeddings were first L2-normalized as

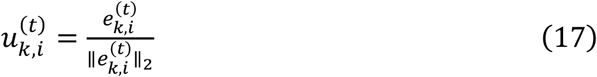

##### Global alignment (*GA*)

measured the similarity between the two participants’ candidate word sets at the level of overall semantic space. It was computed as the mean cosine similarity across all cross-participant word embedding pairs:

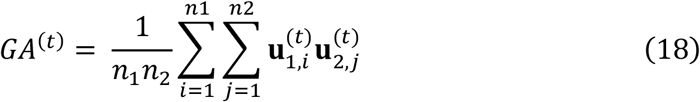

Higher *GA* values indicate that the two participants’ candidate word sets occupy more similar regions of semantic space.

##### Local alignment (*LA*)

measured the similarity between the two participants’ lexical selection distributions within their shared candidate space. To compute this measure, the two participants’ candidate sets were merged into a union set, with overlapping candidate words represented only once. Let **p**^(t)^and **q**^(t)^denote the two participants’ selection probability distributions over this merged candidate space, after zero-padding for absent items and normalization to sum to 1. Local alignment was defined as the negative Jensen–Shannon divergence between these two distributions:

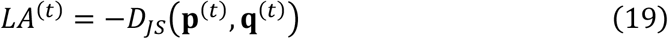

where

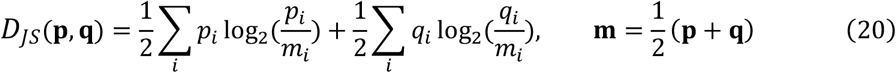

Because lower Jensen–Shannon divergence indicates greater similarity between the two distributions, taking its negative value allowed larger *LA* values to indicate stronger local alignment.

##### Global uncertainty (*GU*)

measured how dispersed a participant’s candidate word set was in semantic space. For each participant, *GU* was then defined as the mean cosine distance across all distinct embedding pairs of candidate words within the set:

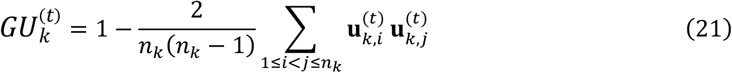

Higher *GU* values indicate that the candidate words are more dispersed in semantic space and therefore reflect greater uncertainty at the global level.

##### Local uncertainty (*LU*)

measured uncertainty in selecting a specific word within the candidate space. For each participant, let **p**^(t)^ = (*p*_1_, …, *p*_*n*_) denote the posterior word-selection probability distribution over *n* candidate words. Entropy was computed as

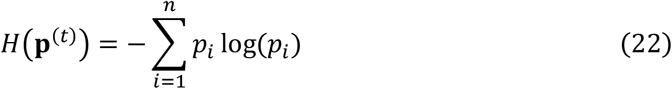

and normalized by the logarithm of the number of candidate words *n*:

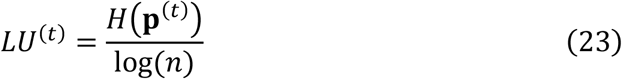

Higher *LU* values indicate greater uncertainty in selecting a specific word from the available candidates.

All four measures—*GA,LA,GU*, and *LU*—were first computed at the trial level. To examine their temporal dynamics, trial positions within each round were converted to percentile positions along the interaction sequence, and the resulting values were averaged across the four rounds of the same task for each dyad or participant. LMEs were fitted with aligned trial position as a fixed effect for each measure in the *Exact communication* task. For task-specific analyses, LMEs included Task, aligned trial ID, and their interaction as fixed effects, with dyad and participant identity as the random intercepts.

#### Prediction of lexical alignment from hypothesis-landscape dynamics

To examine whether dynamic updating of the SPH-derived hypothesis landscape predicted lexical alignment, we quantified trial-to-trial changes in global and local uncertainty. For each participant, condition, and adjacent trial pair, changes in both indices were computed from trial (t-1) to trial (t). Because lexical alignment was measured at the dyadic level, participant-level global and local changes were averaged across partners within each dyad for each trial transition. These dyad-level global and local changes were then used to predict lexical alignment at the later trial. Prediction analyses were conducted separately for each condition using leave-one-dyad-out cross-validation. In each fold, a two-predictor LASSO regression model, including both global and local uncertainty, was trained on all dyads except one and evaluated on the held-out dyad. Prediction accuracy was computed within each held-out dyad as the Pearson correlation between observed and predicted lexical alignment across trials. The model’s overall prediction performance was then defined as the mean of these dyad-level correlations.

Predictor-specific contributions were assessed by comparing the full model with shuffled-feature baselines. In the all-shuffled baseline, both predictors were shuffled; in the selective-shuffling models, one predictor was preserved while the other was shuffled. Shuffling was performed only in the training data and repeated 1,000 times. The full model was directly tested against the all-shuffled and selective-shuffling baselines, with empirical one-tailed *p* values computed from the proportion of repetitions in which it failed to outperform the baseline.

To test whether alignment in the two subcomponents of the SPH-derived hypothesis landscape predicted lexical alignment, we repeated the leave-one-dyad-out prediction analysis using two dyad-level predictors: global and local alignment. The training-set-only shuffling and permutation-testing procedures were otherwise identical to those described above.

#### fNIRS data collection

Two independent NIRsport2 systems (NIRx Medical Technologies, NY, USA) were employed, with each system measuring one individual in a dyad using three customized sets of 44 measurement channels (CHs; Fig. 1b). Two sets of probes were used to cover the bilateral frontal, temporal, and parietal cortices, and third set of probes was used to cover the prefrontal cortex. The anatomical location of each channel was determined by the international 10-20 system. Specifically, CH21, CH43, and CH23 were placed on T7, T8, and Fpz, respectively. The position of the probe sets was checked and adjusted before the experiment to ensure consistency between the 2 communicators in a dyad and among dyads.

To confirm the anatomical locations of the optode probes, we obtained magnetic resonance imaging (MRI) data from 2 female and 2 male participants (not involved in the fNIRS study but drawn from the same population). These participants wore plastic caps on which the probes’ true positions had been marked with Vitamin E balls. They were scanned using a high-resolution, T1-weighted, magnetization-prepared, and rapid gradient-echo sequence (repetition time = 2,530 ms; echo time = 3.30 ms; flip angle=7°; slice thickness=1.3 mm; in-plane resolution=1.3×1.0 mm^2^; and number of interleaved sagittal slices=128). Statistical Parametric Mapping 12 (Welcome Department of Cognitive Neurology, London, UK) was used to normalize the MRI data to the standard Montreal Neurological Institute (MNI) coordinate space^68^. The MNI coordinates of probes were generated according to the Automated Anatomical Labelling template^69^ using the NIRS-SPM toolbox^70^ (see Table S17). Based on this information, we were able to check the consistency between the probes’ true positions and the expected anatomical positions in order to adjust the probes’ true positions. This procedure was repeated several times until the true positions and the expected positions reached a high level of consistency, e.g., the difference of probabilities between the true and expected positions in localizing a specific brain region was <10% for 3 out of 4 scanned participants.

The optical density of near-infrared light (760 and 850 nm) was measured at a sampling rate of 10.17 Hz. Then, changes in oxyhemoglobin (HbO), deoxy-hemoglobin, and total hemoglobin concentration were calculated based on the modified Beer– Lambert law. Here, we focused only on HbO concentrations because HbO is a sensitive indicator of the change in regional cerebral blood flow and has a high signal-to-noise ratio^71^.

#### fNIRS data preprocessing

To ensure the quality of fNIRS data, a running-window procedure was used to calculate the percentage of motion artifacts^72,73^. Channels with >10% of data points identified as containing motion artifacts were labeled as bad and excluded, and participants were excluded if >10% of their channels were bad. Based on these criteria, no participants were removed. The mean percentage of artifacts across all channels and participants was 1.765%±1.484% (mean±SD) in the word generation task and 1.494%±1.453% (mean±SD) in the localizer task.

Next, homer3 functions^74^ were used to preprocess the data. Specifically, motion artifacts were detected and corrected using the discrete wavelet transformation filter procedure^75^, and global physiological noises (e.g., skin blood flow) were removed by using principal component analysis with an 80% variance threshold^76^. Finally, a task-specific band-pass filter was used to remove high- and low-frequency noises in the collaboration word generation task. Specifically, the frequency band of interest (FOI) ranged from 0.038 Hz to 0.195 Hz, corresponding to the period between 5.133 seconds and 26.161 seconds that represents the median of the minimum and maximum trial durations (from listening to the end of speaking) across participants in the word generation task. For the localizer task, the frequency range was set to 0.01–0.5 Hz^72,77^

#### Representational similarity analysis

For the RSA analyses, we first constructed a SPH-based hypothesis landscape RDM for each participant by computing the cosine distance between every pair of posterior-probability–weighted word embeddings (hypothesis landscape) across *Exact communication* trials. The posterior probabilities were derived from the SPH model using the optimal parameters, and the word embeddings were obtained from the GPT-2 model. The corresponding brain RDM was constructed by computing the Euclidean distance between pairs of averaged HbO signals at each channel during the fifth thinking window, where the functional connectivity effect peaked in the preceding analysis (Supplementary Text 6.2). We then calculated the Spearman correlation between the SPH-based hypothesis landscape RDM and the brain RDM for each participant and averaged these correlations across participants. Additionally, to test whether participants also represented their partner’s word-selection process, we constructed the SPH-based hypothesis landscape RDM using the partner’s posterior-probability-weighted word embeddings and again correlated this RDM with the brain RDM. Statistical significance of the RSA correlations was assessed using permutation tests (1,000 iterations). In each iteration, we applied a random permutation to both the rows and columns of the SPH-based hypothesis landscape RDM (self or partner), while keeping the brain RDM fixed, and recomputed the Spearman correlation for each participant and channel. These permuted correlations were then averaged across participants to yield a null distribution of RSA values for each channel. One-tailed empirical *p* values were obtained as (*k*+1)/(1000+1), where *k* is the number of permuted mean correlations greater than the observed mean correlation. Finally, *p* values were corrected for multiple comparisons across channels using FDR.

Next, to test whether the SPH-based hypothesis landscape representation was specific to the *Exact communication* task, we repeated the RSA for two control conditions using the same procedure. For each participant and channel, we first computed the RSA correlation in the *Exact communication* task and the mean correlation across the two control tasks; their difference (*Exact communication* minus baseline) yielded a participant-level RSA correlation difference, which was then averaged across participants to obtain the observed RSA correlation difference for each channel. We then assessed the significance of this difference using a permutation-based test. Specifically, for each task condition, we generated 1,000 permutations by randomly permuting the rows and columns of the SPH-based hypothesis landscape RDM (while keeping the brain RDM fixed). For each permutation, we computed participant-level RSA correlations for the two control conditions, averaged them to yield a permuted baseline, and subtracted this baseline from the permuted *Exact communication* correlations. These permuted RSA correlation differences were then averaged across participants to form a null distribution of RSA correlation differences for each channel. The empirical *p* value for each channel was obtained by comparing the observed RSA correlation difference with this null distribution, and *p* values were subsequently FDR-corrected across channels.

#### Replication of the above main findings based on a different NLP model (FastText)

To replicate the main findings using an NLP model with less pre-trained context information, we replicated main findings in main text using the FastText model. Unlike GPT-2, which is optimized for next-word prediction and sequence generation, FastText is primarily designed for word representation tasks. As such, the candidate word sets originally generated by GPT-2 were retained for consistency. All observed words and candidate words were converted into vector representations (300 dimensions) using the Chinese version of the FastText model (https://fasttext.cc/docs/en/crawl-vectors.html). Other analytical procedures mirrored those applied in the GPT-2-based analysis. Replication results are reported in the Supplementary Text 7.

#### Replication of the above main findings in an independent participant sample and dataset

To further validate our results, we replicated the main findings using a publicly available dataset comprising 20 dyads^43^ (data available at https://ars.els-cdn.com/content/image/1-s2.0-S1053811920311824-mmc2.xlsx). In this dataset, a similar *Exact communication* task and a control task were conducted in Czech, but the words had been analyzed after translation into English in the original and this study. The English version of the GPT-2 model (12 layers, https://huggingface.co/openai-community/gpt2) was employed to generate the candidate word sets and vectorized all target and candidate words in this dataset (768 dimensions). Additionally, to minimize lexical redundancy, all English words were lemmatized to their base forms using the spaCy library in Python. Subsequently, word frequency and part-of-speech (POS) annotations were derived from the Brown corpus, accessed via the Natural Language Toolkit (NLTK) package in Python. All other analytical procedures were identical to those applied to the Chinese dataset. The replication results are presented in the Supplementary Text 8.

### Statistical analysis

Unless otherwise specified, hierarchically structured data (e.g., participants nested within interacting dyads and repeated trial-level observations from the same participant) were analyzed using LME models. Random effects for participant and/or dyad were included as appropriate, and degrees of freedom were estimated using the Satterthwaite approximation. Data without such a hierarchical structure (e.g., independent dyad-level measures) were analyzed using repeated-measures ANOVAs or paired-sample *t* tests, as appropriate. Multiple comparisons were corrected using the Benjamini–Hochberg false discovery rate (FDR) procedure.

## Data and code availability

All analyses were performed using Matlab (R2020b), Python (3.9.1) and R (4.3.0) with standard functions and toolboxes. The data and custom analysis code used in this study will be made publicly available upon peer-reviewed publication.

## Acknowledgments

We thank X. He and B. Liu for their assistance with data collection, F. Zhou for sharing the narrative story material, and other members of L. Zhu’s and C. Lu’s labs for extensive discussions. This work was supported by the National Natural Science Foundation of China (62293550, 62293551) and Interdisciplinary Research Foundation for Doctoral Candidates of Beijing Normal University (BNUXKJC2412).

## Author Contributions

T.Z. and C.L. conceptualized and designed the study. T.Z. conducted the experiments, performed the data analyses, and prepared the visualizations. L.Z. provided critical advice on modeling. T.Z. drafted the initial manuscript, and D.J. Shaw, S.Z., L.Z., and C.L. revised and edited it. Project supervision was provided by C.L. and L.Z., and funding was acquired by T.Z. and C.L.

## Declaration of interests

The authors declare no competing interests.

## Notes

### Competing Interest Statement

The authors have declared no competing interest.

